# Neural correlates of ketamine-induced dilation of temporal expectation

**DOI:** 10.64898/2026.10.12.758574

**Authors:** Jack R. Waters, Brian Madeira, Michael N. Shadlen, S. Shushruth

**Affiliations:** Center for the Neural Basis of Cognition, University of Pittsburgh and Carnegie Mellon University, Pittsburgh, PA, USA; Medical Scientist Training Program, University of Pittsburgh, Pittsburgh, PA, USA; Neuroscience Institute, Carnegie Mellon University, Pittsburgh, PA, USA; Zuckerman Mind Brain Behavior Institute, Columbia University, New York, NY, USA; Howard Hughes Medical Institute, Columbia University, New York, NY, USA; Kavli Institute for Brain Science, Columbia University, New York, NY, USA; Department of Neuroscience, University of Pittsburgh, Pittsburgh, PA, USA

## Abstract

Time estimation is critical to many cognitive operations, including decision-making, working memory, and action planning. Neural correlates of interval timing have been described across multiple cortical and subcortical networks, both at a single-neuron level (as ramping activity) and population level (as dynamical motifs). Low-dose ketamine, a dissociative anesthetic and potent antidepressant, is known to distort time perception, but how it affects these neural correlates is unknown. In two male macaques performing a time-anticipation task, we administered subanesthetic doses of ketamine under a pharmacokinetically guided intravenous infusion protocol. We recorded neural activity from the parietal area LIP, a region associated with interval timing. Behaviorally, ketamine caused monkeys to respond as though less time had elapsed than had actually passed under objective clock time, a dose-dependent dilation of time estimation. This behavioral change was accompanied by parallel effects on the neural representation of elapsed time. At the single-neuron level, ketamine flattened anticipatory ramping activity. At the population level, it induced a rotation of delay-period activity in neural state space, away from the pre-ketamine configuration. By linking a graded pharmacological manipulation to concurrent shifts in behavior and cortical neural activity, we show that ketamine dilates perceived time while reshaping LIP population dynamics. These findings offer a route to probe how cortical activity supports time estimation and point to a network mechanism that could contribute to ketamine’s dissociative and antidepressant effects.

## Introduction

The ability to internally track elapsed time is fundamental to behavior. Decision-making (Finnerty et al., 2015), working memory (Coelho et al., 2016), learning (Eichenbaum, 2014; El Haj and Kapogiannis, 2016), communication (Long et al., 2016), and motor planning (Balasubramaniam et al., 2021) all depend on tracking elapsed time, estimating the probability of upcoming events and generating responses at the appropriate time (Paton and Buonomano, 2018; Janssen and Shadlen, 2005). Neural correlates of elapsed time have been described across the brain, including subcortical regions such as basal ganglia (Chiba et al., 2015; Gouvêa et al., 2015) and cerebellum (Ivry and Spencer, 2004; Garcia-Garcia et al., 2024), and in areas of association cortex, such as prefrontal cortex (Bakhurin et al., 2017; Brody et al., 2003; Genovesio et al., 2006; Murakami et al., 2017), medial frontal cortex (Remington et al., 2018b), and posterior parietal cortex (Leon and Shadlen, 2003; Janssen and Shadlen, 2005; Maimon and Assad, 2006; Jazayeri and Shadlen, 2015).

Neural correlates of timing in association cortex have been described at both single neuron and population levels. At the single neuron level, spiking activity ramps up as the organism tracks time, with the slope of the ramp scaling with the expected length of the interval (Narayanan, 2016; Leon and Shadlen, 2003). The pattern of ramping activity can also follow the hazard function, the conditional probability that the awaited event will occur, given that it has not yet occurred (Janssen and Shadlen, 2005). At the population level, ramping activity has been considered as a low dimensional manifestation of a distributed representation of time (Finnerty et al., 2015; Cueva et al., 2020; Stine and Jazayeri, 2025). This perspective proposes that time is represented through coordinated dynamics distributed across the neural population (Zhou et al., 2020). Although timing-related activity has been perturbed locally in a small number of areas (Long and Fee, 2008; Monteiro et al., 2023; Yang et al., 2025), the relationship between distributed cortical population dynamics and the behavioral readout of time perception has not been probed by direct perturbation.

Ketamine, a dissociative anesthetic and antidepressant, offers a way to perturb these timing signals as it is known to reliably distort the subjective sense of time in both humans and animals. Healthy volunteers given subanesthetic ketamine report that time slows, and psychophysical testing shows impaired temporal discrimination (Coull et al., 2011). Monkeys show a parallel effect of a reduction in temporal expectation during anticipation tasks (Brulé et al., 2021; Ameqrane et al., 2015). However, the neural basis of this time distortion has not been established. While ketamine acts on several receptor systems (Johnson et al., 2015), its best-characterized action is antagonism of NMDA receptors, whose currents support persistent activity during working memory and decision-making (Wang et al., 2013; Suda and Uka, 2022). This raises the possibility that ketamine distorts time perception by reducing the slope of ramping activity (Brulé et al., 2021). But the effects of ketamine on cortical networks are complex (Cichon et al., 2023; Roussy et al., 2021), and its consequences for the population neural dynamics that represent time are unknown.

We set out to determine how ketamine alters the neural representation of time at both the single-neuron and population levels, and whether these changes track the distortion of perceived time. We trained two macaque monkeys on a time-anticipation task, in which they learned to anticipate the go-cue for a memory-guided saccade. We delivered subanesthetic ketamine by continuous intravenous infusion, using a pharmacokinetic model-based dosage regimen designed to hold the plasma concentration of ketamine steady across blocks of trials. Under ketamine, the animals behaved as if less time had elapsed than objectively had, consistent with a dilation of their internal estimate of time. As a window into the distributed cortical networks that track time, we recorded single units from area LIP of parietal cortex, an area known to carry representations of elapsed time (Janssen and Shadlen, 2005; Leon and Shadlen, 2003; Jazayeri and Shadlen, 2015). At the single-neuron level, ketamine reduced the slope of ramping activity, concordant with the observed effect of behavioral time dilation. Ketamine also altered activity of neurons that did not ramp and this manifested at the population level as a rotation of delay period activity away from the pre-ketamine configuration. Together, these results connect ketamine’s dilation of perceived time to changes in cortical activity at both single neuron and population levels. They also provide insight into how ketamine reshapes cortical network dynamics in ways that could contribute to its dissociative and antidepressant effects.

## Results

We trained two male rhesus macaques (Monkey Ch and Monkey Yo) on a memory-guided saccade task. On each trial, the monkey remembered the location of a briefly flashed peripheral target and made a saccade to that location when cued (Figure 1A). The delay between the target offset and the go-cue (’*delay duration*’) varied from trial to trial according to a Weibull distribution (Equation 1, Figure 1B, dashed black curve). The monkeys were rewarded for choosing the correct location with a liquid reward. The reward magnitude was inversely proportional to reaction time (RT), incentivizing the animals to anticipate the go-cue.

**Figure 1:**
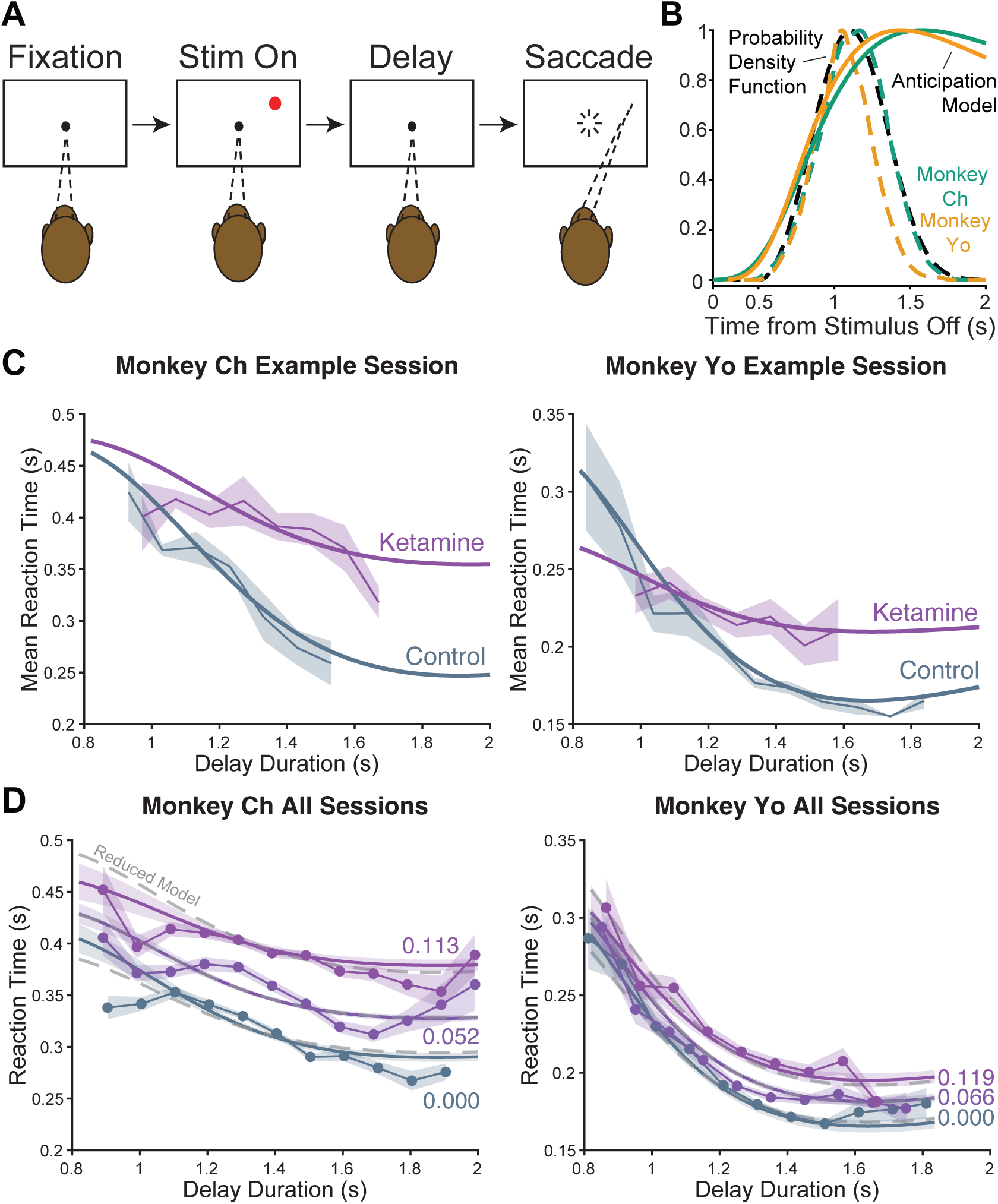
Ketamine dilates estimated elapsed time. **(A)** *Task.* On each trial, the monkey fixated a central spot while a target was briefly flashed at a peripheral location. After a variable delay, disappearance of the fixation point instructed an eye movement to the erstwhile target location. Liquid reward was delivered in a quantity inversely proportional to reaction time (RT). **(B)** *Anticipation model.* Dashed curves show the delay distributions scaled to their peaks: the generative Weibull distribution (black) and the distributions experienced by each monkey (colored). Solid curves are the monkey-specific subjective anticipation models derived from the respective experienced distributions (Equation 3). **(C)** *Ketamine flattens anticipation.* Mean RT (±SE) as a function of delay duration for an example session from each monkey, under control (gray) and steady-state ketamine (purple) conditions. Solid curves are fits to the anticipation models in *(B)*. **(D)** *Concentration-dependent effect of ketamine on anticipation.* Gamma GLM fits (Equation 6) at three representative plasma ketamine concentrations (mg/L) overlaid on mean RTs for the corresponding trials, pooled across sessions. The full range of plasma concentrations overlaid on individual trials is shown in Supp. Fig. S2A. Gray dotted lines show fits to a reduced model without an interaction term.

After training, both monkeys learned to anticipate the timing of the go-cue. The RTs decreased as the delay duration increased, reflecting the animals’ knowledge of the delay distribution (example sessions in Figure 1C, gray). This anticipatory behavior can be described by a subjective hazard function of the delay distribution (Janssen and Shadlen, 2005) (Equation 3, see Methods). This model of anticipation (Figure 1B, solid lines) was estimated separately for the two monkeys based on the delay distributions they each had experienced across sessions (Figure 1B, colored dashed lines) and provided a good description of the RT data for each animal (Figure 1C, gray).

### Ketamine dilates internal estimate of elapsed time

To examine the effect of ketamine on anticipatory time estimation, we administered the drug through an indwelling intravenous catheter in a gently restrained arm while monkeys performed the task. In an initial pilot session, we identified for each monkey the minimal subanesthetic ketamine dose that produced a measurable change in anticipatory RTs. In subsequent sessions, we employed a pharmacokinetically guided infusion regimen to gradually reach and maintain target plasma ketamine levels long enough to characterize its effects on both behavior and neural activity (example session in Supp. Fig. S1, see Methods).

At the steady-state ketamine level, both monkeys showed a flattening of their anticipation functions (example sessions in Figure 1C). RTs during control and steady-state ketamine were comparable at shorter delay durations, but diverged progressively at longer delays, with slower RTs under ketamine. This reduced anticipation of later go-cues is consistent with previous reports that ketamine blunts temporal expectation in monkeys (Brulé et al., 2021; Ameqrane et al., 2015).

To quantify this effect across sessions, we fit RTs with a gamma generalized linear model (Equation 6). The model expressed log RT in terms of the value of the anticipation function at the time of go-cue, the trial-wise plasma ketamine concentration, and their interaction (Figure 1D, Supp. Fig. S2A). In both monkeys, the model with the interaction term accounted for behavior significantly better than a model without it (Monkey Ch: *F* (1, 5327) = 14.5, *p* < 10^−3^; Monkey Yo: *F* (1, 3068) = 4.64*, p* = 0.03; nested F-test; see *Reduced model* fits in Figure 1D). The estimated delay-dependent effect of ketamine was positive in both monkeys (*β*_3_ = 1.38 in Monkey Ch and 1.21 in Monkey Yo), i.e., increasing ketamine concentration reduced the slope of the anticipation function. In contrast, the delay-independent effect of ketamine on RTs was inconsistent across animals (Ch: *β*_2_ = 0.99, *p* < 10^−3^, Yo: *β*_2_ = 0.16, *p* = 0.72). The delay-dependent slowing indicates that, for a given objective delay, the animals behaved as though less time had elapsed — a dilation of subjective time. The RT effect closely tracked the plasma ketamine concentration across the session (Supp. Fig. S2B). And following the cessation of ketamine infusion, the anticipation function returned to its pre-ketamine form in both monkeys (Supp. Fig. S2C).

To test whether the delay-dependent effects of ketamine reflected a generalized deficit in task performance, rather than a change in time estimation, we examined several behavioral metrics (Supp. Fig. S3A). Both monkeys continued to make saccades to the appropriate target location with high accuracy in both control and ketamine conditions (91.8% vs. 95.5% in Monkey Ch and 84.4% vs. 85.6% in Monkey Yo), indicating that the memory of the target location remained intact. Other task outcomes were consistent with a dilation of estimated time. Premature saccades decreased in both monkeys as expected if ketamine led them to under-anticipate the go-cue (Monkey Ch: -1.68%, *p* < 10^−5^, Monkey Yo: -3.97%, *p* < 10^−4^, one-versus-rest mixed-effects model). In Monkey Ch, this led to an increase in the fraction of rewarded trials (+4.5%, *p* < 10^−3^, binomial mixed-effects model). However, in Monkey Yo, the same reduction in premature saccades was outweighed by an increase in trials with no response within the reward window, resulting in a decrease in the rate of rewarded trials (−14.9%, *p* < 10^−22^). The timing of no-response trials in this monkey was telling. They occurred preferentially at shorter go-cue delays (Supp. Fig. S3B). Such a failure to respond to cues that occurred early in the delay is what is expected from a dilated internal clock since the go-cue arrives before the animal expects it. If the animal had forgotten the target location or otherwise disengaged, the no-responses would preferentially increase later in the trial.

The effect of ketamine on trial aborts was in opposite directions in the two monkeys, decreasing in Monkey Ch (-1.68%, *p* = 0.016) and increasing in Monkey Yo (+2.63%, *p* < 10^−3^). Since aborts arise from blinks and fixation breaks, this across-monkey difference likely reflects ketamine’s effect on oculomotor behavior in the two animals, which cannot account for ketamine’s effect on the RT-anticipation interaction. Together, the behavioral effects support a slowing of the internal estimate of elapsed time rather than a generalized memory loss or disengagement.

We next asked whether ketamine’s effects on saccade execution could account for the delay-dependent RT effect. Saccade endpoint error increased slightly but significantly (+0.37 *deg*, *p* < 10^−5^ in Monkey Ch; +0.58 *deg*, *p* < 10^−19^ in Monkey Yo; mixed effects model, Supp. Fig. S4B), indicating a small reduction in motor precision, though the endpoints remained within the acceptance window and thus did not affect overall accuracy. Peak saccade speed also decreased in both Monkey Ch (−7.39%, 95% CI [8.28, 6.50], *p* < 10^−52^, mixed effects model) and Monkey Yo (−13.9%, 95% CI[ 15.06, 12.72], *p* < 10^−92^, Supp. Fig. S4C). However, a uniform change in saccade vigor or precision should affect the delay-independent term (*β*2), and not the interaction term (*β*3). To confirm this, we directly added saccade speed and endpoint distance to the gamma GLM (Equation 10). The interaction remained significant in both animals (Monkey Ch: *β*_3_ = 1.52, *p* < 10^−4^, Monkey Yo: *β*_3_ = 1.24*, p* = 0.005, Supp. Fig. S4D). Thus, the delay dependent effect of ketamine on RT reflects an effect on time estimation rather than a generalized motor deficit.

### LIP single neurons show heterogeneous responses during the delay epoch

To investigate the neural basis of the behavioral time dilation induced by ketamine, we recorded spiking activity from 210 isolated single units (126 from Monkey Ch, 84 from Monkey Yo) from area LIP of posterior parietal cortex. Recordings were targeted to parts of ventral LIP that showed strong persistent activity during memory-guided saccades (Figure 2A–B, Gnadt and Andersen (1988)). A prior study demonstrated that neurons in this area represent the hazard rate of elapsed time (Janssen and Shadlen, 2005) on trials where the target was shown in their response field (RF). That study showed that delay-period firing rate of many single neurons increases as a function of the hazard rate of the anticipation function. Population averages were also reported to reflect this response profile. Our multichannel recordings sampled a larger and more heterogeneous population than the previous study. Individual neurons differed in how they represented elapsed time during the delay period. Some neurons ramped up in anticipation of the go-cue whereas other neurons ramped down (example neurons shown in Figure 2C–D, gray). This diversity suggests that representation of elapsed time could be distributed across neurons with a range of temporal response profiles. The relative proportions of the different response phenotypes differed across the two monkeys, leading to different population average response profiles (Figure 2E–F).

**Figure 2:**
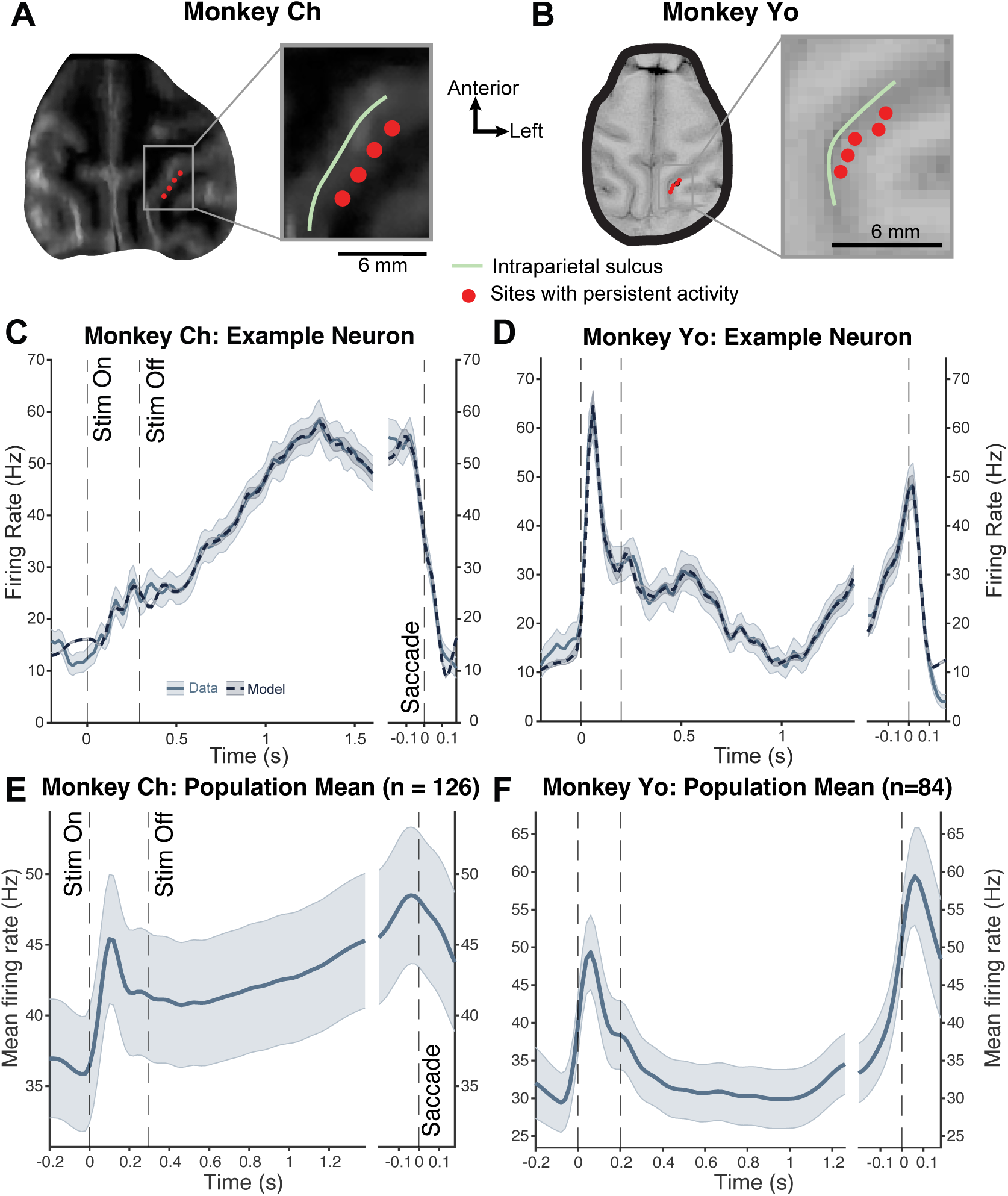
Single neuron and population averages in LIP represent elapsed time. **(A-B)** *Recording sites.* MRI sections for each monkey (oriented orthogonal to the electrode trajectories) showing the locations with persistent delay-period activity (red dots). Recordings were targeted to these sites. Inset shows magnification around the intraparietal sulcus. **(C-D)** *Example neurons.* Peristimulus time histogram (PSTH, mean SE) of two example neurons on trials with the target in the response field. Across the delay period, firing rate increased in one neuron *(C)*, and decreased in the other *(D)*, illustrating the heterogeneity of time representation across neurons. Dashed lines show overlaid fits of a generalized linear model (see Figure 3A). **(E-F)** *Population average.* Average PSTH (mean SE) across the recorded LIP population in each monkey, reflecting the different dominant response profile in each animal.

A source of confound for understanding the diversity of neural responses during the delay period is that it includes the sensory responses to the target flash and saccade initiation motor responses. To isolate the delay-specific neural responses, we used a Generalized Linear Model (GLM) based approach (Park et al., 2014). This approach allowed us to isolate the component of delay-period spiking activity that carries timing signals by decomposing each neuron’s trial-by-trial spiking activity into separable contributions from the sensory response to the target flash, delay-period activity, motor activity around saccade initiation, and spike history (Figure 3A).

**Figure 3:**
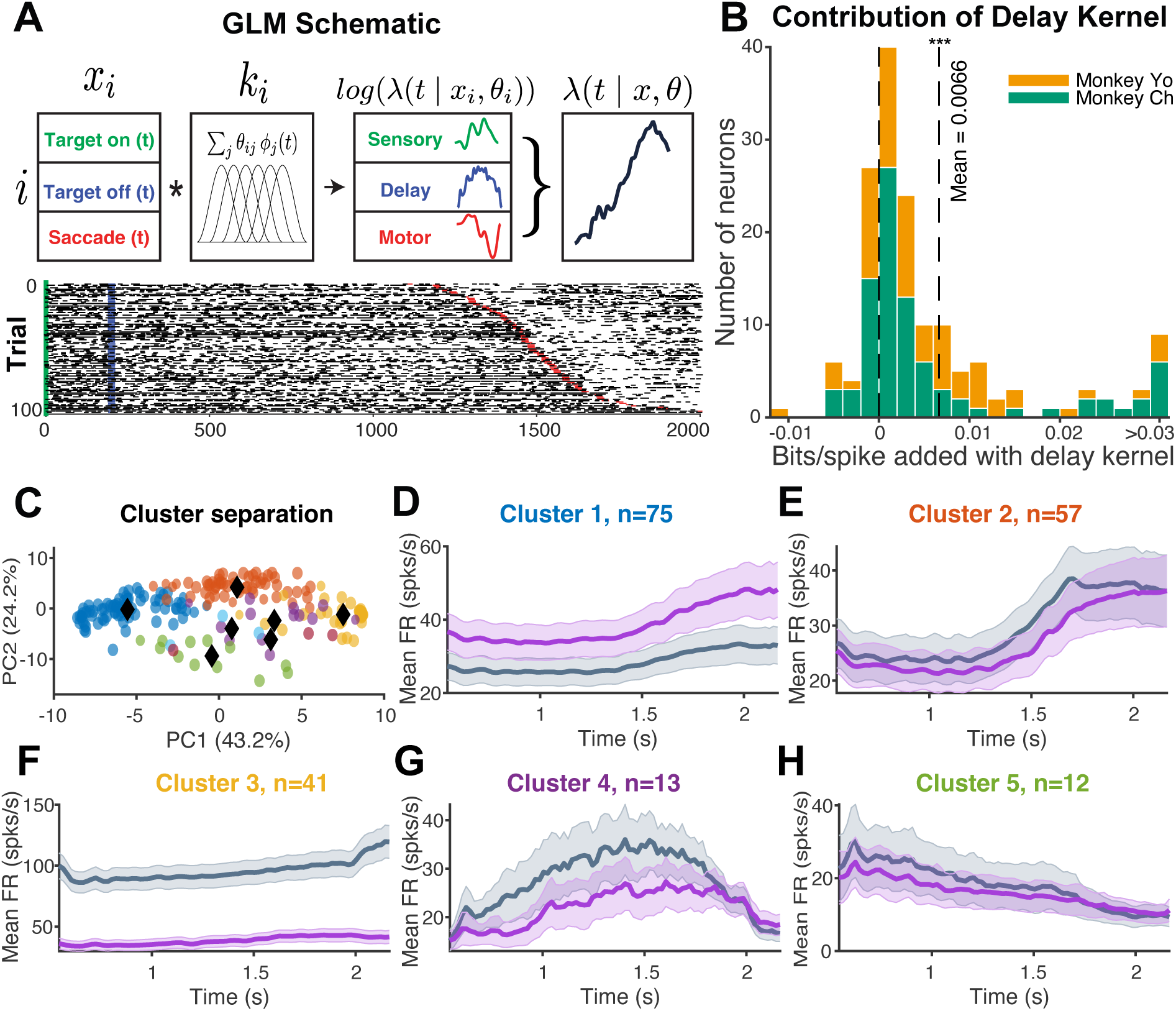
Single neurons show diverse delay-period signals. **(A)** *GLM Schematic.* The generalized linear model decomposes trial-by-trial spiking activity into four components: sensory response, delay-period activity, motor preparation, and spike history. Each component is a temporal kernel (*k_i_*), fit as a weighted sum of raised cosine basis functions (Equation 11). **(B)** *Delay kernel improves spike prediction of control data.* Cross-validated model performance (bits/spike) compared between the full model (with delay kernel) and a reduced model (without delay kernel) for each neuron, in the two monkeys. Values greater than zero indicate neurons for which including the delay kernel improved spike prediction. **(C)** *Cluster separation.* All valid neurons were combined across both monkeys and clustered by their GLM delay kernels using a Gaussian Mixture Model. Clusters are shown in the top two principal components, with black dots marking the cluster centers. **(D-H)** *Cluster profiles.* Mean delay kernel firing rates for the five largest clusters, under baseline (gray) and steady-state ketamine (purple). Each panel’s label color matches the cluster’s location in (*C*). Mean responses of each cluster separated by monkey are shown in Supp. Fig. S5.

We first determined whether each neuron had delay-period modulation beyond those accounted for by either the visual response or motor preparation. To do so, we asked if the inclusion of a separate *delay kernel* in the GLM improves spike prediction on held out trials. For the majority of neurons that met trial count criteria (see Methods), the delay kernel improved this cross-validated spike prediction (67/88 neurons in Monkey Ch and 53/70 neurons in Monkey Yo; *p* < 10^−13^, signed-rank test, Figure 3B). The full model provided a good description of single-neuron firing rates across all task epochs (example fits Figure 2C–D, dashed lines). We then used the fitted delay kernels to recover each neuron’s delay-associated spiking activity for subsequent analyses.

Inspection of the delay kernels confirmed the diversity of delay-period modulations across individual neurons. To characterize this heterogeneity, we used a Gaussian Mixture Model (GMM) to cluster neurons based on their delay kernel activity (Figure 3C, see Methods). The clusters spanned a range of delay-period dynamics (Figure 3D-H), including clusters of neurons whose delay activity ramped up, ramped down or showed a biphasic profile, among others. Similar response profiles were seen in populations from each monkey (Supp. Fig. S5). This establishes that activity is modulated in a variety of ways during the delay period, but not whether such modulation has leverage over the animal’s timing behavior. We return to this question at the population level below.

### Ketamine has bidirectional effects on single neuron firing

We next examined how ketamine affected the delay-period activity of LIP neurons. Its effect on the overall firing rate (FR) was bidirectional, decreasing it in some neurons (examples in Figure 4A,D) and raising it in others (Figure 4B,C; ΔFR: Median: 0.19 Hz, IQR: [-5.11, 2.67] Hz; Supp. Fig. S6). Despite this variability in changes to mean firing rate, ketamine had a consistent effect on the slope of the delay-period activity. The first two example neurons from Figure 4 illustrate this. In the neuron whose activity ramped up during the delay-period, ketamine reduced both the overall firing rate and the slope of the ramp (Figure 4A). In the neuron whose activity ramped down, ketamine had the opposite effect, elevating the rate, yet still attenuating the slope of the ramp (Figure 4B). In both cases, therefore, ketamine reduced the slope of the delay-period response, regardless of its polarity. This flattening held across the sampled population in both monkeys (Monkey Ch: *p* < 10^−12^, Monkey Yo: *p* = 0.02, signed-rank test, Supp. Fig. S7).

**Figure 4:**
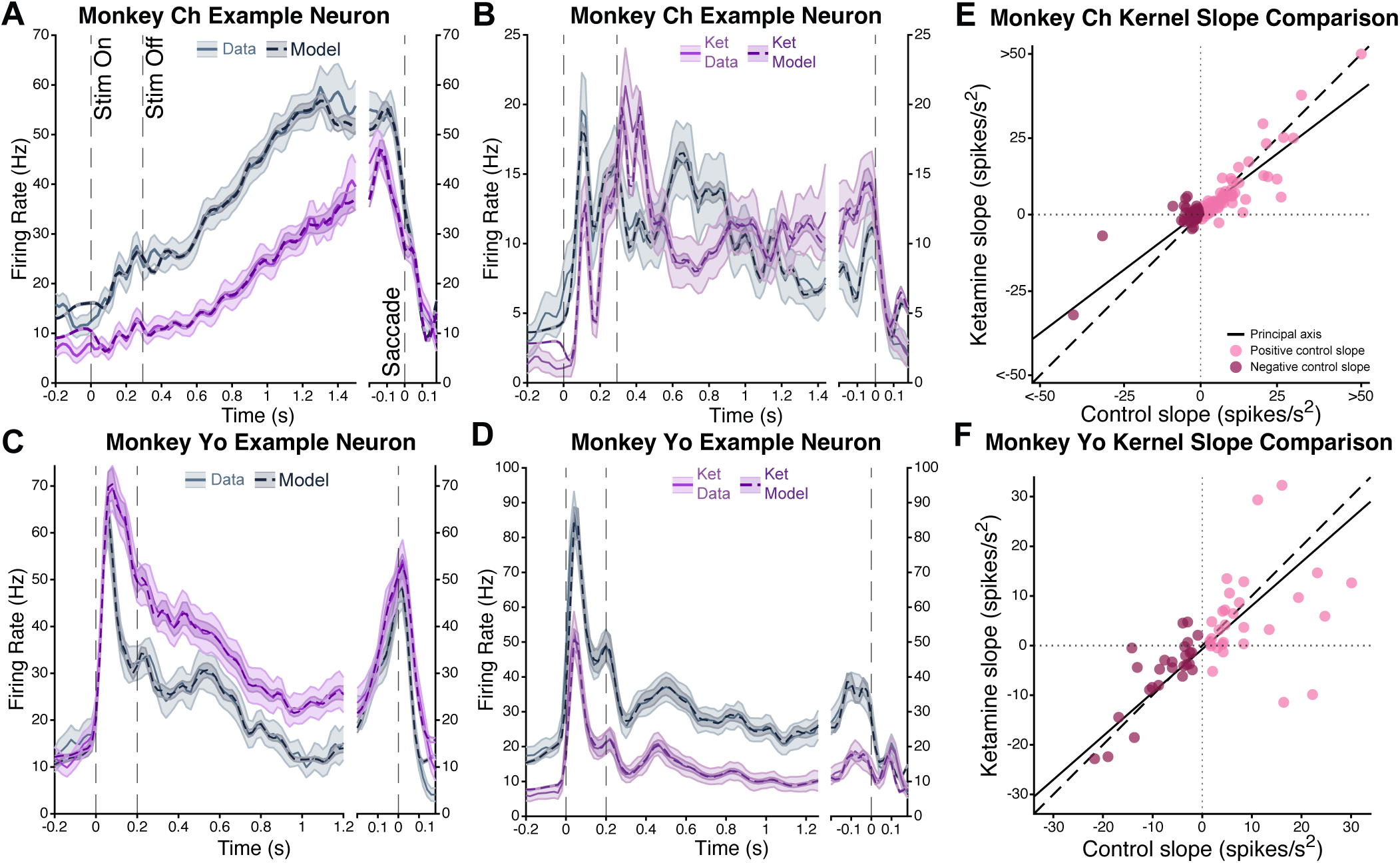
Ketamine flattens ramping slopes. **(A-D)** *Effect of ketamine on the PSTH.* PSTHs of four example neurons under baseline and steady-state ketamine. Dashed lines are GLM fits, estimated separately for control and steady-state ketamine trials. **(E-F)** *Kernel slope comparison.* Delay kernel activity slope under control versus steady-state conditions, plotted separately for each monkey. Points represent individual neurons and are colored by the sign of baseline slope (upward or downward ramping). Principal axis is shown as a solid line and the unity line as dashed.

As mentioned above, the start and end of the raw delay-period responses are susceptible to sensory and motoric influences respectively. We therefore computed the change in slope of the delay kernel activity, which isolates delay-period activity from sensory and motor components. Ketamine again significantly reduced the magnitude of the slope of ramping activity and this result held when separately assessed for upward-ramping (Monkey Ch: *p* = 10^−4^, Monkey Yo: *p* = 0.03, signed-rank test) and downward ramping neurons (Monkey Ch: *p* < 10^−5^, Monkey Yo: *p* = 0.01, Figure 4E-F). Together, these results show that ketamine reduces the anticipatory ramping activity during the delay period, consistent with the behaviorally observed dilation of internal time estimates.

The effect on slope is part of a broader diversity of ketamine effects across the clusters of delay-period modulation (Figure 3D-H). Although we defined the clusters from the control delay kernel activity, they differed in how they responded to ketamine as firing rates rose in some clusters and fell in others. Thus the slope reduction during the delay period that correlates with behavioral time dilation (Figure 4E-F) is likely part of a more complex effect.

To characterize the effect of ketamine on single neurons more systematically, we incorporated plasma ketamine concentration as a predictor into the GLM (Equation 13, see Methods). This allowed inclusion of trials at intermediate ketamine concentrations, thus increasing the number of neurons with sufficient trials for model fitting. This ketamine-parameterized model predicted held-out spike trains above the homogeneous Poisson baseline across the 191 neurons included in this analysis (mean 10-fold cross-validated performance: 0.21 bits/spike above baseline; median: 0.14 bits/spike). For a vast majority of these neurons (102/111 in Monkey Ch and 79/80 neurons in Monkey Yo), inclusion of a delay kernel in the ketamine-parameterized model improved spike prediction (Signed-rank test: median relative gain in bits/spike 2.6%, *p* < 10^−29^, Supp. Fig. S8), indicating that ketamine reshaped delay-period activity rather than only scaling the overall rate. The model also permitted predictions of neuronal responses for a range of ketamine plasma concentrations, revealing a graded relationship between delay-period firing dynamics and plasma concentration (example neurons in Supp. Fig. S9). These dose-dependent effects were heterogeneous across neurons and over the course of the delay period. We next address how these varied single-neuron changes combine to affect the population-level representation of elapsed time.

### Ketamine reshapes behaviorally relevant population dynamics

The delay-period modulations described so far establish that LIP activity changes over the delay, but not that these changes bear on the animal’s timing behavior. We therefore asked whether delay-period population activity carries information about the monkeys’ trial-by-trial anticipation. We utilized demixed principal component analysis (dPCA, Kobak et al. (2016)) to isolate a population dimension that captured the interaction between delay duration and RT, i.e., whether the monkey responded faster or slower than predicted for a given delay. In the pre-ketamine condition, projections onto this dimension separated slow from fast trials (Figure 5A), so we refer to this dimension as the “*behavioral time axis*”. This dimension explained significantly more unique population variance than expected following permutation of fast and slow trial labels (*p* = 0.004, permutation test). This behavioral time axis continued to explain significant unique variance when it was estimated after removing neurons from the two upward ramping clusters (Figure 3D-E), although the effect was weaker (*p* = 0.018, permutation test, Figure 5B). Moreover, removing any single cluster before fitting, including the upward and downward ramping clusters, did not reduce the unique variance the axis explained relative to a size-matched null, demonstrating that no single cluster exerted a uniquely significant influence on the behavioral time axis (Supp. Fig. S10). Behaviorally-relevant timing information was therefore distributed across the sampled neural population and not carried by ramping neurons alone.

**Figure 5:**
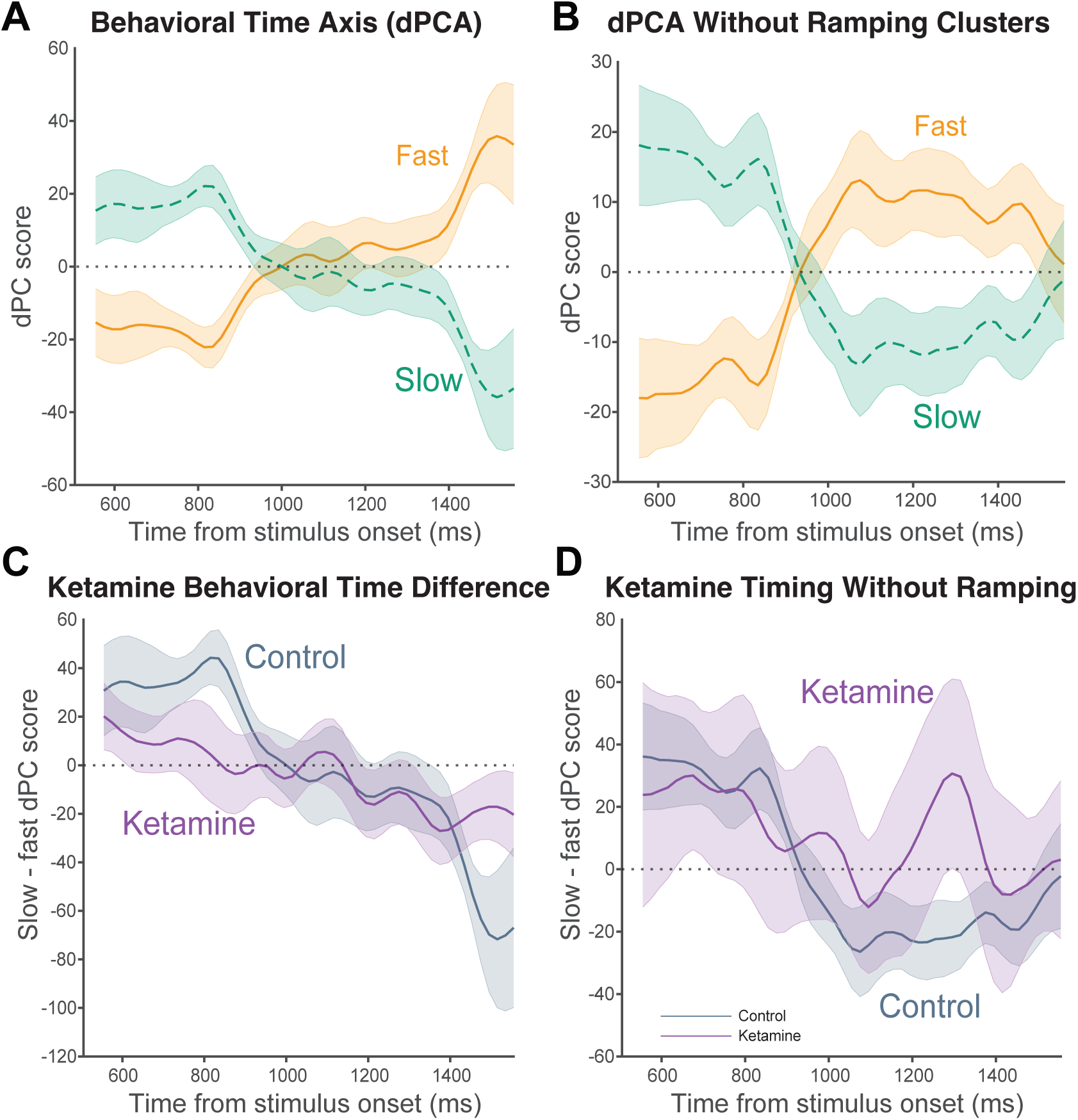
A behaviorally relevant population activity axis. **(A-B)** *Behavioral time axis.* Projection of control neural activity (±SE from bootstrap) onto an axis identified by dPCA that captures time-dependent differences between fast and slow RT trials, which we term the ’*behavioral time axis*’. In *(A)*, the axis was determined using all neurons; in *(B)*, the axis was determined without the largest upward ramping clusters (*D* and *E* in Figure 3). Because the axis captures the difference between fast and slow trials, the two projections are approximately antisymmetric about zero. **(C-D)** *Ketamine’s effect on the behavioral time axis.* Separation between the fast and slow trial projections along the behavioral time axis is shown for control (gray) and steady-state ketamine (purple). *C* and *D* use the behavioral time axes from *A* and *B*, respectively. The separation was reduced under ketamine, but the reduction was not significant.

Having identified a population axis that tracks the animal’s timing behavior, we asked how ketamine changes activity along it. We projected the trials at steady-state ketamine condition onto the behavioral time axis. The separation between slow and fast trials was smaller under ketamine, but this reduction was not significant (Figure 5C, *p* = 0.3, permutation test). The same trend was present, again not significant, when the analysis was restricted to non-ramping neurons (Figure 5D, *p* = 0.22). Thus, while the direction of the effect of ketamine on the behavioral time axis is consistent with the single neuron and behavioral effects, the size of the effect was too small to reliably establish at the population level. Two factors might have contributed: the limited number of trials at steady-state ketamine, and the fact that dPCA analysis was applied to raw firing rates during the delay period which also carry sensory and motor responses. So we turned to the GLM-derived delay firing rates, which isolate delay-period modulations across ketamine concentrations.

### Ketamine rotates the delay-period subspace while conserving trajectory shape

The population representation of timing has been described as a trajectory through neural state space (Remington et al., 2018a). In this view, populations represent different intervals by varying the speed with which the neural state evolves along a fixed trajectory (Sohn et al., 2019; Remington et al., 2018a). If ketamine acted this way to dilate temporal expectation, the neural dynamics should evolve slower under ketamine (Hypothesis 1, Figure 6A *H*_1_). An alternative possibility is that ketamine changes the orientation of the population trajectory relative to the control delay-period subspace (Hypothesis 2, *H*_2_), consistent with ketamine’s effect of redistributing activity across neural populations (Cichon et al., 2023). Under *H*_2_, rotation away from the control subspace would shorten the trajectory’s projection onto the control dimensions. This could produce a dilation in behavioral timing if downstream readout remains aligned to those dimensions (Kaufman et al., 2014).

**Figure 6:**
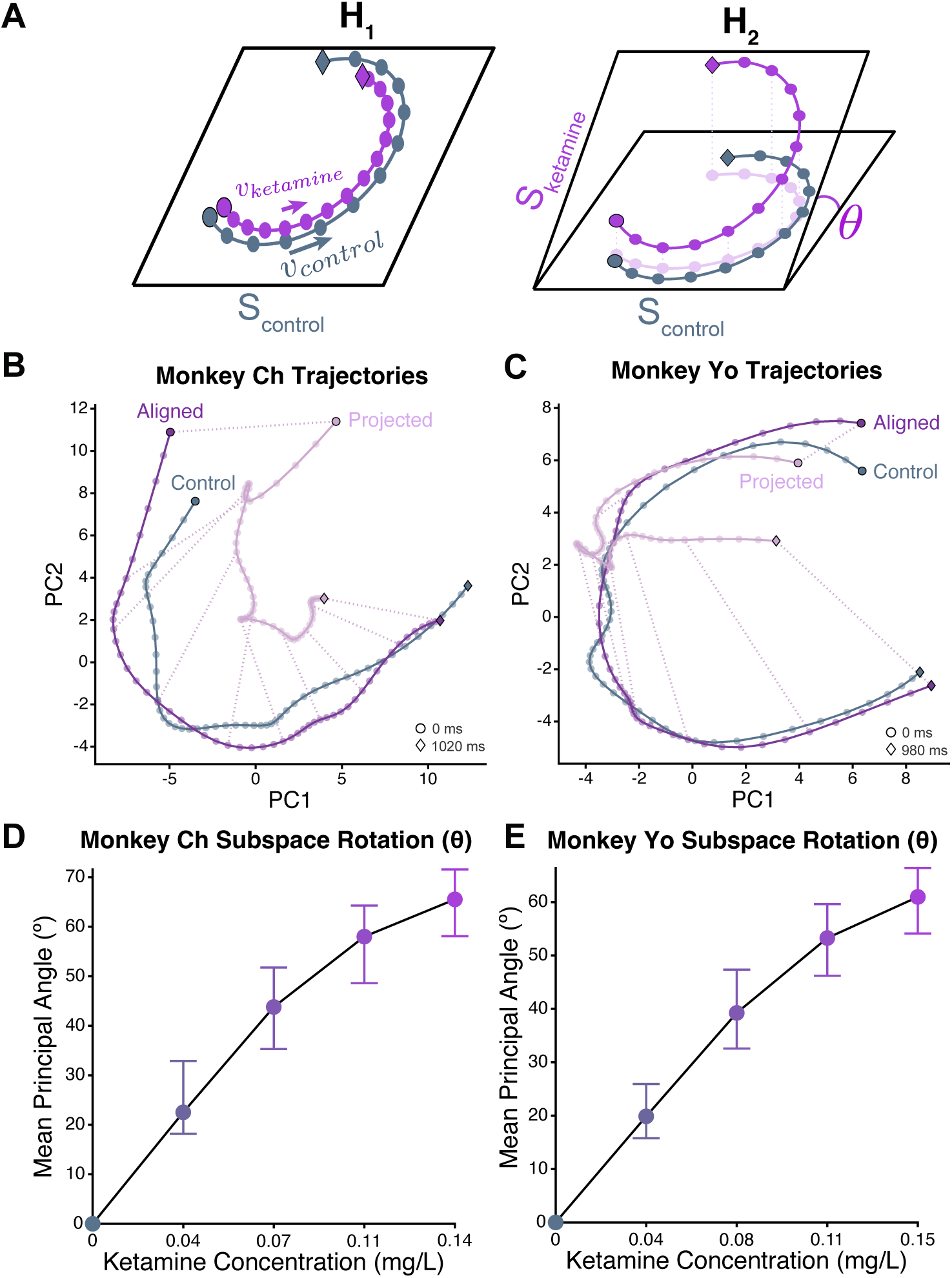
Ketamine induces dose-dependent rotations in population activity. **(A)** *Schematic of hypotheses for changes to trajectories under ketamine.* Under H1, ketamine reduces the speed along the same trajectory in the control subspace *S_control_*. Under H2, ketamine induces a subspace rotation by an angle *θ*. **(B-C)** *State space trajectories.* Delay kernel activity for control (gray) and steady-state ketamine (pink) data are projected onto the top two principal components of the control subspace (*S_control_*). Both trajectories span the same duration, with circles marking the start and diamonds the end of the delay-period kernels. The same ketamine activity from its own subspace (*S_ketamine_*) is also shown aligned to the control trajectory using Procrustes alignment without scaling (purple). Dashed lines connect matched time points on the aligned and projected ketamine trajectories. Aligned trajectories at other doses are shown in Supp. Fig. S11G-H. **(D-E)** *Subspace rotation.* Mean principal angle between *S_control_* and *S_ketamine_* (with bootstrapped 95% CIs) is shown for a range of ketamine plasma concentrations. The angle increases progressively with dose in both monkeys.

To test these hypotheses, we applied principal component analysis (PCA) to the GLM-derived delay kernel activity across the population of neurons that were significantly delay modulated, i.e., those for which the delay kernel improved cross-validated prediction (Supp. Fig. S8). Each neuron contributed a single delay kernel, and we defined the control subspace (*S_control_*) from the control kernel activity of these neurons. In both monkeys, most of the variance in the control data was captured by a small number of dimensions (92.7% in 3 dimensions in monkey Ch and 93.1% over 4 dimensions in Monkey Yo). The neural trajectories in this low dimensional space (*S_control_*) evolved systematically over the course of the delay interval (Figure 6B-C, gray). This established a putative population correlate of time in the pre-ketamine control condition, similar to previous studies (Meirhaeghe et al., 2021).

We then projected delay kernel activity for the same set of neurons across a range of ketamine plasma concentrations onto *S_control_* (projection of the highest concentration shown in pink in Figure 6B-C). These projections decreased in length with increasing ketamine plasma concentration (Supp. Fig. S11A-B). We confirmed that this result held for trajectories computed from raw firing rates, instead of the delay kernel activity (Supp. Fig. S11C-D). The shortening of trajectory lengths is expected under both *H*_1_ and *H*_2_. To adjudicate between these two hypotheses, we asked whether ketamine induced a rotation in population dynamics away from the dominant control delay-period subspace. For each ketamine concentration, we computed the dominant subspace, *S_ketamine_* from the corresponding delay kernel activity and measured the mean principal angle between *S_control_* and *S_ketamine_*. The angle progressively increased by approximately 4.45° per 0.01 mg/L increase in plasma ketamine for Monkey Ch (95% CI: [3.09, 4.69]) and 4.05° for Monkey Yo (95% CI: [2.76, 4.23]) (Figure 6D-E). We ruled out the possibility that changes across principal angles were driven by changes to minor dimensions by assessing angle as a function of explained variance (Supp. Fig. S11E-F). This rotation accounts for the shortening of the projected trajectories in *S_control_*, since a trajectory tilted out of a subspace leads to a shorter projection.

Finally, we asked whether ketamine only tilted the dominant subspace or also reshaped the dynamics within it. We aligned the control and steady-state ketamine trajectories with Procrustes analysis, allowing rotation, reflection, and translation but not scaling (Equation 15). We compared the trajectory shapes by computing an *R*^2^ metric between the aligned trajectories (Equation 16). The trajectory shape was well conserved in both monkeys at the maximal ketamine concentration (Figure 6B-C, dark purple) and across a range of intermediate concentrations (Supp. Fig. S11G-H, Monkey Ch *R*^2^: mean = 0.94; range = [0.89, 0.98]; Monkey Yo *R*^2^: mean = 0.95; range = [0.92, 0.97]). Ketamine therefore caused a systematic, dose-dependent reorientation of delay-period dynamics while maintaining their shape.

## Discussion

Time estimation is a fundamental computation distributed across the brain, and supported by diverse neural correlates (Finnerty et al., 2015; Stine and Jazayeri, 2025). However, this ubiquity poses a challenge for causally linking these neural correlates to an organism’s sense of time. Ketamine, an anesthetic and antidepressant that is known to distort time perception (Coull et al., 2011), provided an ideal tool to approach this problem. Because the effect of ketamine on neural representations of time was also unknown, the same experiments allowed us to characterize these properties of a clinically important drug. Consistent with previous behavioral studies (Brulé et al., 2021; Ameqrane et al., 2015), we found that ketamine dilated the subjective estimate of time i.e., the animals behaved as if their internal clocks ran slow relative to the objective passage of time. By recording from one node of the timing network, parietal area LIP, we found that this behavioral dilation was accompanied by concordant changes at both single-neuron and population levels. We first consider the specificity of the behavioral effects of ketamine, then what the perturbation reveals about the neural correlates of timing, and end with what it implies for ketamine’s effects on cortical networks.

### Targeting time estimation with ketamine

Our task required the monkey to track elapsed time continuously, and it provided a readout of the monkey’s sense of time on individual trials through RTs as a function of delay duration. While this readout enabled us to quantify ketamine’s graded effects on timing, it also posed a problem since ketamine is an anesthetic and slows movement. To separate the two effects, we delivered the drug via pharmacokinetically guided intravenous infusion, which let us slowly reach and maintain a low, subanesthetic concentration. In contrast to many previous studies that used a single bolus dose, our approach allowed us to isolate the effects of ketamine on timing in detail.

Three features of the animals’ behavior indicate that the observed effects are predominantly specific to timing. First, in both monkeys, the effect of ketamine on RT depended on delay duration. This was supported by the significant interaction between ketamine dose and delay duration, in addition to the main effect of the drug. Ketamine thus changed how RT varied with delay, and not simply the overall time to respond. Second, the animals continued to perform the task accurately. Saccades continued to be directed to the appropriate target, indicating that the memory of the target location was unaffected and the animals remained engaged in the task. Third, while the drug had some effect on the saccade itself, these changes were found to be insufficient in explaining the delay-dependent effects of ketamine. Thus, the slowing of RTs does not simply reflect a general deficit in saccade execution. Together, these results establish that our pharmacokinetic approach isolated a predominantly timing-specific effect of ketamine, which allowed us to ask how cortical signals of timing were affected.

### Perturbing the neural correlates of timing

Ramping of firing rate is a canonical single-neuron correlate of interval timing. The slope of the ramp reflects the length of the interval being timed, and the firing rate in each instant indexes how much of that interval has elapsed (Narayanan, 2016; Leon and Shadlen, 2003; Jazayeri and Shadlen, 2015). Persistently active neurons can also carry richer information about elapsed time, such as the hazard of an awaited event – its conditional probability of occurrence, given that it has not yet occurred (Janssen and Shadlen, 2005). With the advent of larger population recordings during timing tasks, a diversity of population activity patterns are now observable. Progression through these patterns as an interval elapses is now commonly described as a trajectory through neural state space (Remington et al., 2018a). These single neuron and population level descriptions are complementary, and both covary with timing behavior (Stine and Jazayeri, 2025).

These neural correlates have been described across many cortical (Genovesio et al., 2006; Remington et al., 2018a; Leon and Shadlen, 2003) and subcortical areas (Gouvêa et al., 2015; Ivry and Spencer, 2004; Yang et al., 2025), suggesting a globally distributed network for time tracking. This ubiquity complicates causal investigation of the link between these correlates and behavior with local manipulations, as perturbations of activity in one area can be compensated by modulation of activity in another (Jeurissen et al., 2022). Thus, we leveraged the behavioral effects of systemically administered ketamine as a tool for causal manipulation. The goal of this manipulation is to understand the effects on the neural correlates of timing themselves rather than on the individual areas that carry them. In this spirit, the recordings from area LIP are meant to provide insight into the effects of ketamine on cortical timing networks, rather than on what happens in a particular brain region.

The neural effects of ketamine were consistent with behavioral time dilation at both the single-neuron and population levels. Ketamine reduced the slope of ramping activity, as has been hypothesized (Brulé et al., 2021), but not previously demonstrated. Importantly, slopes were flattened for both up-ramping and down-ramping neurons in a fashion not accounted for by a simple suppression of firing rate. Ketamine therefore modulated patterns of time-related signals, rather than simply suppressing activity.

This single-neuron effect foreshadowed a change in the population as a whole, as captured by two measures. First, ketamine reduced the separation along dimensions of population activity that best discriminated the animals’ timing behavior, i.e. fast vs. slow reaction times. This is the population signature most directly tied to behavior, and ketamine appeared to distort it, although this reduction did not reach significance. Second, population trajectories within the dominant control subspace were shorter over the same duration with higher concentrations of ketamine. This effect is consistent with a decrease in speed, as would be expected from the results of previous research linking behavioral timing to population dynamics (Remington et al., 2018b; Sohn et al., 2019; Meirhaeghe et al., 2021; Stine and Jazayeri, 2025), but it was only true when dynamics were projected into the control subspace. In contrast, when population trajectories were instead expressed in their own dedicated coordinate spaces, we found that trajectory shapes were highly conserved. Each of these measures are most easily explained by a rotation of activity away from the dominant subspace, indicating that ketamine causes a redistribution of activity across neurons.

One potential mechanism by which a rotation in population activity could produce a change in time estimation is by altering the relationship between population representations of elapsed time and their readout for behavior. Under control conditions, timing-related activity might occupy a behaviorally relevant subspace aligned with downstream circuits which integrate these signals across areas to drive timed action. Reorientation of activity away from these output dimensions may reduce its influence, and thereby alter the rate of integration. Providing credence to this idea, work in rodents suggests that cortical timing signals might be integrated downstream in basal ganglia (Yang et al., 2025).

### Insights into ketamine’s effects on cortical networks

We have shown that ketamine’s effect of dilating behavioral time estimation is associated with a rotation in population dynamics. Rotations across subspaces of neural activity are an area of increasing interest for their mechanistic role in a number of neural computations, including preparing for motor action (Kaufman et al., 2014; Churchland and Shenoy, 2024) and maintaining information in working memory (Libby and Buschman, 2021), highlighting the ability of cortical networks to redistribute representations to achieve desired outputs. Our results extend this framework by demonstrating that rotations can also occur as a result of pharmacological perturbation with ketamine.

This raises a question of how ketamine might produce such a rotation. Although our investigation centered on the effects of ketamine on timing, this does not imply that its effects are restricted only to this domain. Rather, we posit that ketamine affects a general property of cortical networks that timing depends upon.

One candidate is the balance between excitation and inhibition (Shadlen and Newsome, 1994; van Vreeswijk and Sompolinsky, 1996). Excitatory-inhibitory (E-I) balance shapes recurrent network dynamics, including their intrinsic timescales (Wang, 2002; Murray et al., 2014), and therefore could alter both the evolution and orientation of population dynamics. Previous work indicates that low-dose ketamine alters E-I balance (Cavanagh et al., 2020), which leads us to propose that ketamine modulates interactions between neurons, reorienting the dominant timing subspace.

Consistent with this view, ketamine’s effects on single neurons are heterogeneous, raising the firing rate in some and decreasing it in others. This effect of redistributing excitation and inhibition across the population has been previously documented, and reflects the complex receptor pharmacology of the drug (Johnson et al., 2015). While we were unable to examine the role of specific cell types in driving this rotation, ketamine is thought to most strongly affect parvalbumin- and somatostatin-expressing inhibitory neurons (Cichon et al., 2023; Roussy et al., 2021), which may suggest a key role for these cell types in driving rotations during other, unperturbed neural computations.

Irrespective of the mechanism, characterizing the effects of ketamine at the network level has wider implications. It can unify the effects of low dose ketamine on other cognitive operations, such as working memory (Corrigan et al., 2025; Blackman et al., 2013). Further, it bears on the clinically important aspects of the drug, including its use as a dissociative anesthetic and an antidepressant. A holistic account of how ketamine works would require tying together its actions at the level of receptors and network dynamics to behavior. Whole-brain models built at cellular resolution (Schmidt et al., 2018) offer a route to that account. Our results are a step towards that goal.

## Acknowledgements

We thank the veterinary staff of Columbia University, in particular Dr. Catrina King and Amanda Blanchard, for help with ketamine administrations. We thank Faisal Baqai for help with data processing and spike sorting and Kendra Noneman for providing code to process eye signals. We thank Drs. Diana Burk, Steve Chase, Danique Jeurissen, Jon Johnson, Caroline Runyan, Matthew Smith, and Saurabh Vyas for helpful discussions, and Dr. Yasmine el-Shamayleh, Akhil Bandi and Ezra Sutter for comments on an earlier version of the manuscript. This work was supported by HHMI (MNS), a University of Pittsburgh Medical Center CMRF Award (SS) and computing resources from the University of Pittsburgh Center for Research Computing (HTC cluster supported by NIH award S10OD028483).

## Methods

### Experimental Model and Subject Details

All training, surgery, and experimental procedures were conducted in accordance with the Public Health Service Policy on Humane Care and Use of Laboratory Animals (National Research Council, 2011). Experiments were approved by the Columbia University Institutional Animal Care and Use Committee (IACUC) under protocol number AC-AAAW4454. We trained two adult male rhesus macaques (*Macaca mulatta*) – Monkey Ch (age 17 years, weight 12 kg) and Monkey Yo (age 18 years, weight 10.5 kg) – on a time-anticipation task and recorded neural activity from parietal area LIP.

### Behavioral Task

Monkeys performed a memory-guided saccade task (Figure 1A). The animal initiated a trial by fixating on a central red point. After 0.6–1 s of maintained fixation (drawn from a truncated exponential distribution), a target appeared. The target location was drawn from 4-15 positions, chosen each session to match the response fields of the recorded neurons. The target was presented for 200 ms (Monkey Ch) or 300 ms (Monkey Yo), followed by a memory delay drawn from a Weibull distribution of the following form, where *α* = 3, *τ* = 0.5:

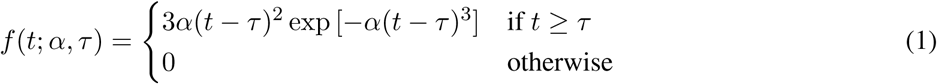

Extinguishing the fixation point served as the go-cue, instructing the monkey to saccade to the remembered target location for a juice reward. To encourage anticipation of the go-cue, reward size decreased with saccade latency. The reward magnitude function was set separately for each monkey and adjusted across sessions.

Visual stimuli were presented on a CRT monitor (75 Hz refresh rate; viewing distance 48 cm). Eye position was recorded using an infrared eye tracker (Eyelink, SR Research; sampling rate: 1 kHz). Stimuli were generated using either Expo software (Peter Lennie, New York University) for Monkey Ch, or the Psychophysics Toolbox (Kleiner et al., 2007) in Matlab (Mathworks) under the control of ReX experimental control system (Hays Jr et al., 1982) for Monkey Yo. Juice rewards were delivered by a solenoid-based system.

### Neural Recordings

Each monkey was implanted with a headpost for head fixation and a CILUX recording chamber (Crist Instruments) over the left parietal cortex under general anesthesia and aseptic conditions. We used magnetic resonance imaging (MRI) to localize the intraparietal sulcus (IPS) in relation to the recording chamber. We obtained MR images (inversion recovery fast spin echo in Monkey Ch and T1 weighted gradient echo in Monkey Yo) with a recording grid *in situ* and used custom software to project the recording grid onto the MR images. In pilot recording sessions, we systematically mapped the lateral bank of the IPS (sampling every 300–500 *µm*) and noted the locations of neurons with spatially selective persistent activity (Figure 2A-B) during visually-guided and memory-guided saccade tasks (Gnadt and Andersen, 1988). Recordings were made with either three independently controlled single electrodes (Minimatrix, Thomas Recordings) or 16-channel V-probes (Plexon Inc). Electrode placement varied across sessions but was targeted to sites on the lateral bank of the IPS with strong delay-period multi-unit neural activity. The raw voltage signal was recorded at 30 kHz (OmniPlex, Plexon Inc.) and single units were extracted using the Wave clus spike sorting algorithm (Chaure et al., 2018).

### Ketamine Administration

Monkeys were trained to perform the task with their arm in a custom built gentle restraint that allowed the placement of an intravenous (IV) catheter that stayed in place throughout the session. An infusion of saline was started through an IV line driven by a syringe pump (Harvard Apparatus Inc.) placed outside the recording rig. After a block of control trials, the infusate was switched to ketamine without any overt cues to the monkey.

In an initial pilot session, we slowly ramped up the ketamine dose to identify the minimal dose that produced a measurable change in anticipatory RTs. We estimated the corresponding plasma concentration with a pharmacokinetic model (see below) and used this concentration as the target for the recording sessions. Using the model, we administered a dosing regimen that slowly achieved the target concentration and maintained it over 10-25 minutes. We conducted 8 and 7 sessions in Monkey Ch and Monkey Yo, respectively.

### Pharmacokinetic Model

We approximated the time course of plasma ketamine during and after infusion with a three-compartment model (Ihmsen et al., 2001; Fanta et al., 2015), integrated in 1-minute steps. The model is composed of three coupled equations corresponding to the changes to ketamine concentrations in plasma (*C*_1_) and two peripheral compartments (*C*_2_, *C*_3_). The concentrations evolve as:

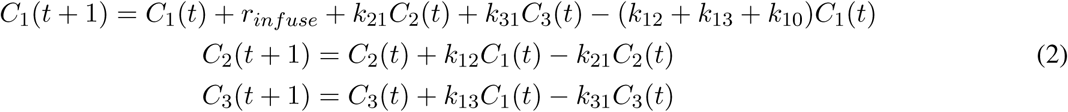

where *r_infuse_* is the intravenous infusion rate of ketamine, and *k*_12_*, k*_21_*, k*_13_*, k*_31_*, k*_10_ are the rate constants relating exchange between the three compartments and elimination from the central compartment. We used rate constants based on human ketamine pharmacokinetics (Ihmsen et al., 2001; Fanta et al., 2015), as there are no published data in macaques. While the exact values likely differ in macaques, they are unlikely to affect the monotonic relationship between infused dose and estimated concentration on which our dose-dependence analyses rely.

Session-specific ketamine infusion dosage regimes were used to estimate plasma concentrations on individual trials (example session in Supp. Fig. S1). We assigned to each trial the estimated concentration at its onset. Trials prior to the first infusion dose were designated as the pre-ketamine control condition, and trials in which the estimated plasma concentration surpassed a threshold concentration of 0.1 mg/L were defined as steady-state ketamine condition.

### Behavioral Data Analysis

We modeled each monkey’s subjective anticipation function following the framework of Janssen and Shadlen (2005). We first constructed the distribution of delay times each monkey experienced using kernel density estimation (normal-approximation method) on correct trials pooled across sessions (Figure 1B, colored dashed lines). Due to variable tolerance for long durations between each monkey, the experienced distribution differs from the generative Weibull (Figure 1B, black), as it reflects each animal’s specific patterns of aborts and false starts. From the experienced distribution, we derived subjective anticipation as a function of the hazard rate of elapsed time as per:

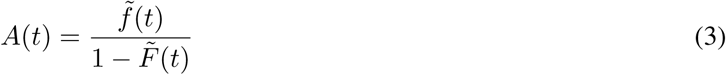

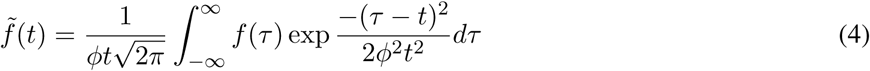

Where *f̃*(*t*) and *F̃*(*t*) are the experienced distribution and its integral, respectively. The convolution with a Gaussian kernel (Equation 4) represents the monkeys’ increasing uncertainty over the course of the delay. The width of the kernel scaled with elapsed time according to a Weber fraction (*ϕ*) of 0.26. Using the experienced rather than the generative distribution ensured that the anticipation function reflected each monkey’s own knowledge of go-cue probability. These subjective anticipation functions provided a good description of RT as a function of delay duration in individual sessions (e.g., Figure 1C-D), when fit parametrically to reaction times (*r*) at a given delay (*t*):

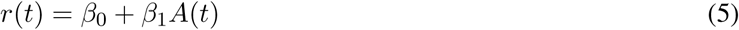

For this initial analysis, we included only trials with a correct saccade and valid reaction time (RT, time from go-cue to saccade) between 150 and 800 ms, excluding missed trials and premature saccades.

### Effect of Ketamine on Behavior

To assess the impact of ketamine dose on RT, we modeled the expected reaction time (*µ_i_*) as a Gamma distribution, as RTs are strictly positive and tend to be right-skewed (Luce, 1986). For each trial, *i*, the log-expectation of RT was linearly related to the value of the subjective anticipation function at the time of go-cue, *A*(*t_i_*), and the trial plasma concentration of ketamine, *K_i_* according to

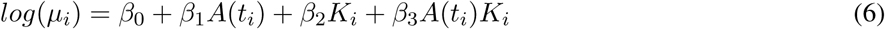

where *β*_1_ and *β*_2_ are the main effects of the subjective anticipation function and the ketamine concentration, respectively, and *β*_3_ is their interaction. Parameters were fit by maximum likelihood estimation using an iteratively reweighted least squares algorithm. Because of behavioral adaptation to the effects of ketamine over the session, trials more than ten minutes following the final infusion of ketamine were excluded from all analyses.

### Eye Movement Analysis

Eye position was continuously recorded at a sampling rate of 1 kHz. Analog eye signals were translated from raw voltage to degrees of visual angle by linearly mapping the median voltages between fixation (300–50 ms prior to target onset) and target acquisition (25–75 ms following acquisition) for correct trials in each session. Calibrated eye traces were then low-pass filtered using a fourth order Butterworth filter with a cutoff of 30 Hz. Saccades were detected from radial eye velocity and acceleration with thresholds of 30 deg./s and 1000 deg./s^2^, respectively. Task-relevant saccades, such as those used to derive reaction time estimates, were additionally required to extend beyond 2° of visual angle to exclude smaller saccades within the fixation window. The first task-relevant saccade within 500 ms of the go-cue was selected and verified with the recorded target acquisition times on each successful trial.

### Analysis of Task Failures

We used eye movement data to evaluate changes to task outcomes at steady-state ketamine plasma concentration. We evaluated four categories of unrewarded trials: misses, premature saccades, aborts, and no-response trials (Supp. Fig. S3A). Misses correspond to trials in which the monkey made a saccade whose endpoint fell outside the acceptance window. Premature saccades occurred before the go-cue. Aborts consisted of blinks and other failures to maintain fixation until the go-cue. If no saccade was detected within 800 ms following the go-cue, this was considered a no-response trial. To test the impact of ketamine on the probability of each category of outcome, we used a one-versus-rest logistic mixed effects model with a fixed effect of condition and a random effect of session:

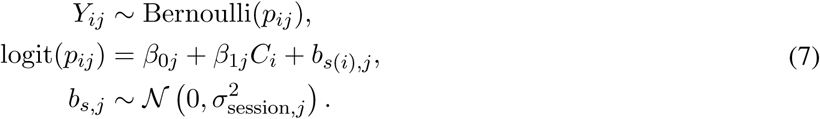

where, *Y_ij_* = 1 when trial *i* has outcome *j*, and 0 otherwise. *C_i_* = 1 for steady-state ketamine and 0 for control. *b_s_*_(_*_i_*_)_*_,j_* is the random intercept for session *s* containing trial *i*. Separate models were fit for the following outcome categories: success, abort, no-response, premature saccade, and misses. Resulting *β*_1_*_j_* values, which represent the log odds of each outcome *j* relative to all other outcomes combined, are reported as percent probabilities using the inverse logit transformation in Supp. Fig. S3A.

Noting a significant change in the probability of no-response trials in Monkey Yo, we then investigated how the monkey’s likelihood of no-response varied with ketamine plasma concentration and delay time (Supp. Fig. S3B). We fit a similar binomial mixed-effects model to Equation 7 using continuous ketamine plasma concentration, delay length, and the delay-ketamine interaction to assess the influence of delay on the probability of no-response:

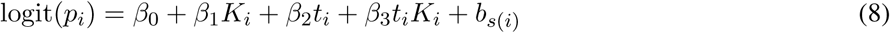

where *K_i_* is the plasma concentration of ketamine and *t_i_* is the delay time for each trial.

### Analysis of saccade metrics

Saccade endpoints were defined as the eye position in the period 0-20 ms following saccade offset or up to the next saccade. Both successful trials and misses were included in the analysis of saccade endpoints, providing a general metric for saccade accuracy. Endpoint error was calculated as the distance from the instructed target on each trial:

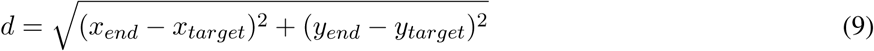

Peak saccade speed was calculated for correct trials in both control and steady-state ketamine conditions. Due to differences in calibration across sessions, both endpoint distance and trial-level log saccade speed were analyzed separately for each monkey using linear mixed effects models with target position and session as random intercepts and ketamine condition as a fixed effect. Model coefficients and confidence intervals are presented as a change in degrees or speed, for each model.

Ketamine had a significant effect on both saccade endpoint error and speed, so we assessed whether the delay-dependent effect of ketamine on RT (Figure 1D) persists when these motor variables were accounted for. We refit the gamma generalized linear model (Equation 6, ’baseline model’) with additional predictors of saccade speed (*v*) and saccade endpoint error (*d*):

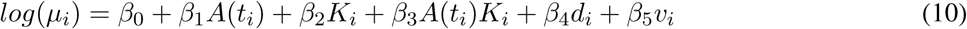

We compared the baseline model to models including either or both of these predictors. All models were fit to completed trials with reliable estimates of endpoint error and speed. We tested whether the ketamine-anticipation interaction (*β*_3_) remains significant once motor effects were included (Supp. Fig. S4D).

### Neural Data Analysis

For each of the 210 isolated neurons, receptive fields (RFs) were defined from peak normalized responses across target locations between stimulus on and off (200-300 ms), on pre-ketamine control trials. All analyses were performed on trials in which the target fell within the RF for a given neuron. The RF borders were defined as the furthest target from the peak response before the sensory response fell below z = −1.5. Unrewarded trials with premature saccades or misses were excluded from analysis. For model fits, trials were additionally excluded if no spikes were observed across the sensory and saccade epochs to avoid matrix singularity or multicollinearity. Neurons with fewer than 20 valid trials were excluded from model fits, resulting in N = 158 for models fit only to control and steady-state ketamine trials, and N = 191 for models that included trials across the full range of plasma concentrations. Peristimulus time histograms were aligned to either stimulus onset or saccade time, computed in 20 ms bins, and smoothed with a 100 ms Gaussian kernel for visualization. Bins following the saccade were omitted from stimulus epoch, bins prior to the saccade were omitted from the saccade epoch, and epochs were divided at the point when saccades were made in 50% of trials.

To examine the effect of ketamine on ramping activity during the delay period, we first identified neurons with a significant ramping slope during pre-ketamine control trials (*p* < 0.05) from linear regression of the firing rates in the time window 100 ms after target offset up to 150 ms before saccade on each trial. For these neurons, we computed the slopes separately for control and steady-state ketamine trials, and compared them with one-sided Wilcoxon signed-rank tests, run separately for upward and downward ramping neurons (Supp. Fig. S7). This analysis was repeated using extracted delay kernel activity from our Poisson generalized linear model (Figure 4E-F), as detailed below.

### Poisson Generalized Linear Model

To isolate delay-associated spiking activity from post-target and pre-saccadic activity at the level of single trials, we utilized a Poisson generalized linear model framework (Figure 3A, Park et al. (2014)). This model predicts the firing rate on single-trials as a linear combination of task events (*x_i_*), convolved with an associated set of kernel weights (*k_i_*). The task events were the target onset (sensory), target offset (delay period), and saccade (motor). Effects of recent spiking were incorporated through a nonlinear post-spike history kernel (*h*), capturing intrinsic spiking dynamics such as bursting, adaptation, and phasic spiking (Weber and Pillow, 2017). History kernel weights were fit using control trials, in the time window -500 ms to -250 ms prior to target onset. Each neuron’s firing rate was assumed to be Poisson distributed with time-varying spike rate *λ* and fit via maximum likelihood estimation. Task event kernels were constructed as a linear sum of overlapping raised cosine temporal basis functions (*ψ_j_*) of 60 ms in width, with temporal weights *θ_ij_*.

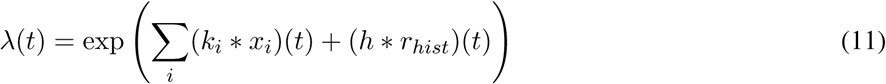

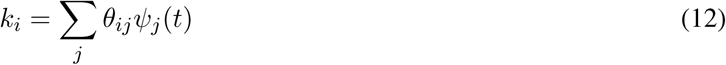

To test whether a neuron carried delay-period activity beyond its sensory and motor responses, we compared the full model to a reduced model without the delay kernel, using cross-validated bits per spike on held-out trials (Figure 3B). This baseline model assumes that periods beyond the sensory and saccade kernels contain Poisson-distributed firing activity. A neuron was counted as delay-modulated if the delay kernel improved held-out prediction. To isolate the delay period activity, the coefficient-weighted sum of delay-kernel bases was exponentiated. The resulting firing rate predictions separate delay period responses from sensory and motor responses.

We used two approaches to characterize ketamine’s effect on firing rate. In the first, we fit two separate models to pre-ketamine control trials and to steady-state ketamine trials, to confirm that the GLM captured ketamine-induced changes in neural responses. Ketamine’s effect on the slope of the delay kernel activity (Figure 4E-F) was analyzed in the same way as the slope analysis on raw firing rate (Supp. Fig. S7). In the second, we directly modeled the effect of ketamine dose on firing rate (Supp. Fig. S9) by incorporating a single estimated ketamine plasma concentration (*K_n_*) for each trial (*n*). Parameterizing ketamine continuously allowed this model to use all trials, including intermediate concentrations excluded from the two-condition fit. Normalized ketamine concentration was included both as a main effect on overall firing rate (*β*_1_) and through an interaction with the delay kernel, allowing ketamine concentration to modulate delay-period timing dynamics as follows:

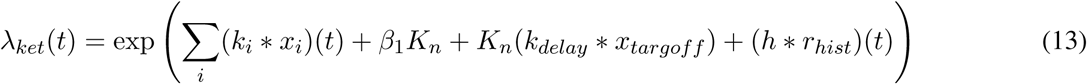

Predictive performance of each model was assessed using 10-fold cross-validation. Folds were stratified to preserve relative numbers of control and ketamine-dosed trials. Performance was quantified using bits per spike, relative to a homogeneous Poisson model whose mean firing rate was estimated from the same training data. Positive bits-per-spike values indicate an improvement in predicted spike trains relative to the constant-rate model. Extraction of ketamine-modulated delay kernel activity consisted of a weighted sum of the delay kernel, the ketamine main effect, and the ketamine-delay interaction kernel, followed by exponentiation.

### Neuron Clustering

To characterize the heterogeneity of delay-period dynamics across our dataset, we utilized a Gaussian Mixture Model (GMM) to cluster all valid neurons by their delay kernel predictions, including those which did not see substantial improvements in information with the addition of the delay kernel (total: N = 198). Utilizing the pre-ketamine control GLM, we extracted kernel-specific predictions for activity over the delay period and z-score normalized across all control and ketamine trials. Neurons were aggregated across both monkeys to identify shared motifs in single unit activity. We used PCA on control kernels to reduce dimensionality across time bins and selected the minimum number of PCs to explain greater than 90% of variance.

Following PCA, a GMM was fit to the control component scores to identify clusters of neurons with similar delay kernel dynamics. The number of clusters was chosen by minimizing the Akaike Information Criterion. Fits with any cluster containing fewer than three neurons were excluded. The mean delay kernel activity for each cluster is shown in Figure 3D-H. The mean of the delay kernel activity from fits to the steady-state ketamine trials are also shown there.

### Behavioral Time Axis

To test whether there was a neural activity dimension that correlated behaviorally with the time dependent RTs, we first divided trials across all recorded neurons according to whether their RT was 15% slower than the predicted reaction time, based on our control Gamma GLM fits to behavior, or else they were categorized as fast trials. We then utilized demixed principal component analysis (dPCA) to identify dimensions of pseudopopulation neural activity which tracked elapsed time and separated fast and slow trials.

Firing rates were first averaged across fast or slow trials for each neuron and separated by ketamine condition. Spike trains were binned in 20 ms increments, smoothed with a Gaussian kernel, and restricted to 200 ms following stimulus offset and 200 ms prior to saccade. dPCA was fit exclusively to pre-ketamine control data and demixed according to reaction time (fast/slow), elapsed time, and the interaction. The leading component of the interaction between elapsed time and reaction time describes how neural activity evolves over the interval for fast versus slow behavioral timing and is thus referred to as the ”behavioral time axis” (Figure 5A-B). Uncertainty was estimated through 1000 bootstrapped samples of neurons, with replacement, and projection bands indicate standard error. Unique axis variance and significance were calculated through a permutation procedure with 1000 repetitions by shuffling fast and slow labels (reaction time and interaction). For permutations of the elapsed time component, each neuron’s time marginalization was circularly shifted by an independently sampled random number of time bins, wrapping values around from the end of the analysis interval to the beginning. This randomized temporal alignment across neurons while preserving each marginalization’s values and cyclic ordering. Unique variance for each axis was estimated by finding the difference in variance explained by the model with and without that axis. To test whether ramping clusters were of unique importance to the behavioral time axis, the same bootstrap and permutation procedures were performed with dPCA fit to the population with upward ramping clusters removed (Figure 3D-E).

### Population Dynamics

To characterize changes to population dynamics under ketamine, we utilized principal component analysis (PCA) to reduce dimensions across our dataset. After simulating a range of ketamine concentrations, we found the trial-averaged firing rate predictions from the delay kernel for each condition (Supp. Fig. S9). Predicted firing rates were binned, and minimal Gaussian smoothing was applied for denoising. We separately verified that this smoothing did not directionally affect any final results. Time-bins were aligned across neurons to stimulus offset. The bins were then truncated at the nearest bin to 1000 ms from target offset, near the mode of the Weibull delay distributions, so that the same duration was analyzed for all neurons, trials, and conditions.

Delay kernels were generated for a range of ketamine plasma concentrations including the control, and neurons were concatenated together to produce matrices of neurons by time bins. Each ketamine concentration matrix was then dimensionally reduced with PCA, and all conditions were projected into the control PCA subspace, determined by the number of dimensions required to achieve greater than 90% explained variance (3 for Monkey Ch, 4 for Monkey Yo). The same analysis was also performed on real firing rates to verify that our results were not restricted to model-based effects (Supp. Fig. S11C-D).

To assess whether differences in control subspace projections were accompanied by changes in the shape of trajectories, we compared these projections to trajectories in their respective condition-specific PCA coordinates (Figure 6B-C, Supp. Fig. S11G-H). For each ketamine concentration condition *k*, we used Procrustes analysis to estimate an orthogonal transformation matrix *R_k_* ∈ ℛ^*d*×*d*^ and translation vector *c_k_* ∈ ℛ^1×*d*^ that aligned the condition trajectory *X_k_* to the control trajectory *Y* . This alignment permitted rotation, translation, and reflection of trajectories without scaling or temporal warping. This procedure minimizes the sum of squared Euclidean distances between the control matrix *Y* and each condition matrix *X_k_*:

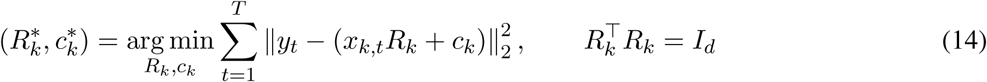

Here, *X_k_*, *Y* ∈ ℛ^*T* × *d*^ contain *T* matched time bins and *d* retained principal components, *x_k,t_* and *y_t_* denote their respective rows, and *I_d_* is the *d*-dimensional identity matrix. The aligned trajectory is given by the following, where 1*_T_* is a column vector of *T* ones:

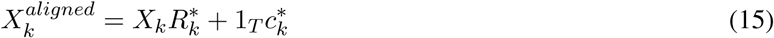

To measure the agreement in trajectory shapes following Procrustes alignment, we calculated *R*^2^ values for each ketamine concentration from the residual sum of squares and total centered sum of squares of the control trajectory:

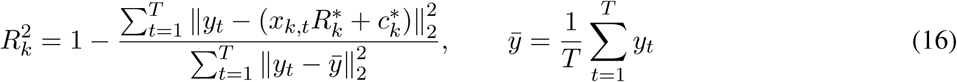

where *ȳ* is the mean position of the control trajectory. Perfect alignment would yield *R*^2^ = 1. As alignment did not permit scaling or warping, this metric reflects agreement in shape, size, and the speed of evolution in dynamics.

Principal angles between subspaces were calculated from PCA loading matrices for each ketamine concentration before alignment (Figure 6D-E, Supp. Fig. S11E-F). Let *Q_Y_*, 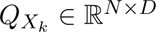 contain the first *D* principal component vectors for control and concentration *k*, where *N* is the number of neurons which demonstrated improvements with the delay kernel in the ketamine-parameterized model (Supp. Fig. S8). To compare directions across subspaces, we used singular value decomposition:

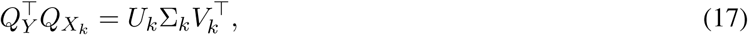

where *U_k_* and *V_k_* are orthogonal matrices containing the left and right singular vectors, respectively, that identify corresponding directions within the two subspaces, and the diagonal entries of Σ*_k_* give the cosines of the angles between these directions. Then, the principal angles are:

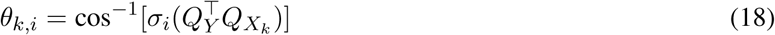

where 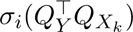 denotes the *i*th largest singular value, or diagonal entry of Σ*_k_*. We summarized these angles using their mean:

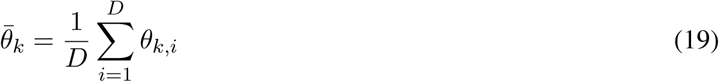

Length of the control trajectory and each projected trajectory was computed as the sum of Euclidean distances between 20 ms bins in PCA space (Supp. Fig. S11A-D). Let *F_k_* R*^N×T^* represent either raw firing rates or the delay kernel predicted firing rates at ketamine concentration *k*. We first projected each condition matrix into the control subspace,

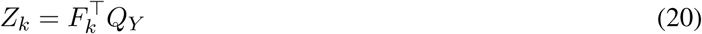

Then length *L* was computed as

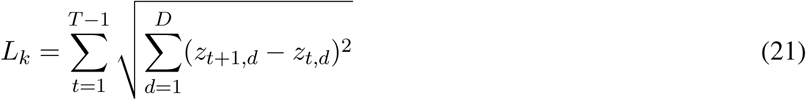

where *z_t,d_* represents the coordinate along the *d*th principal component of the projection matrix *Z_k_* ∈ ℛ^*T* × *D*^ at time bin *t*. Uncertainty in both length and angle measurements was estimated using 1000 paired neuron-bootstrap resamples. The same neurons were sampled with replacement from each condition, and PCA was refit for each sample with dimensions retained up to the resampled control’s 90%-variance dimension. Approximate two-sided p-values for length differences from control were calculated and confidence intervals are presented as the 2.5th and 97.5th bootstrap percentiles.

## Supplemental Figures

**Figure S1:**
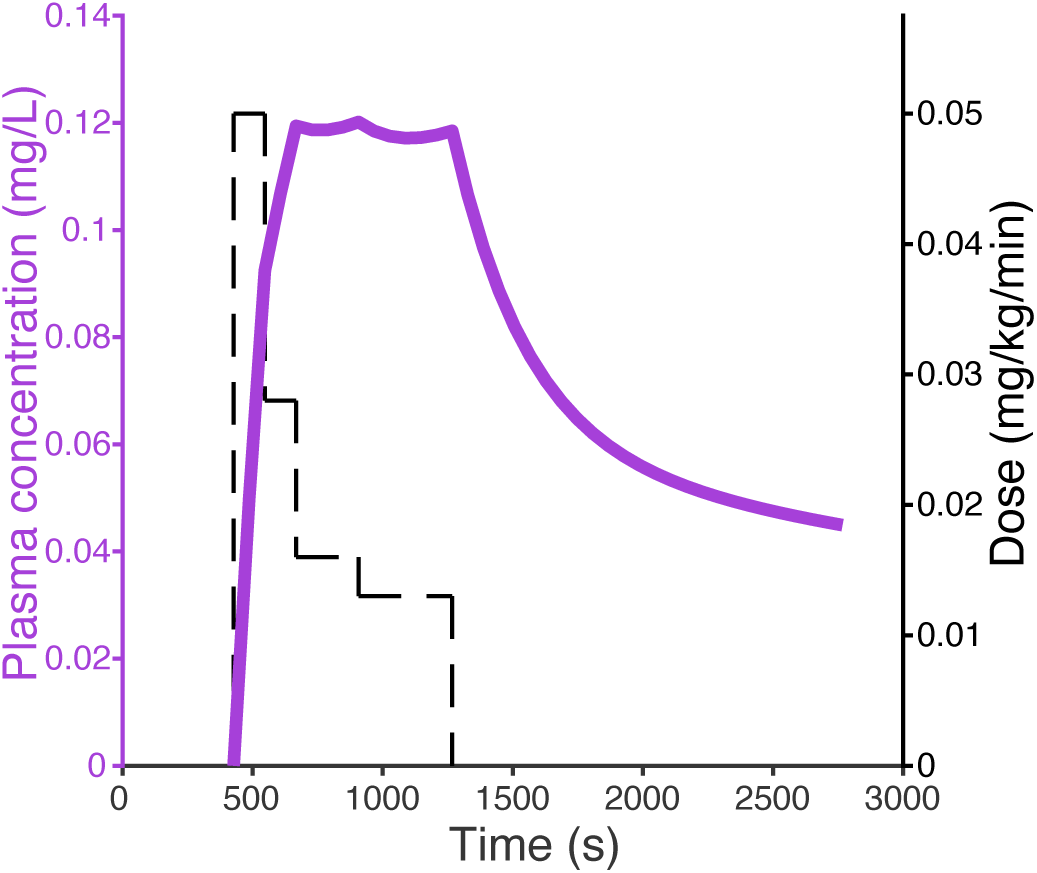
Plasma pharmacokinetics of ketamine. Estimated plasma ketamine concentration over the course of an example session shown with the corresponding dosage of infused ketamine.

**Figure S2:**
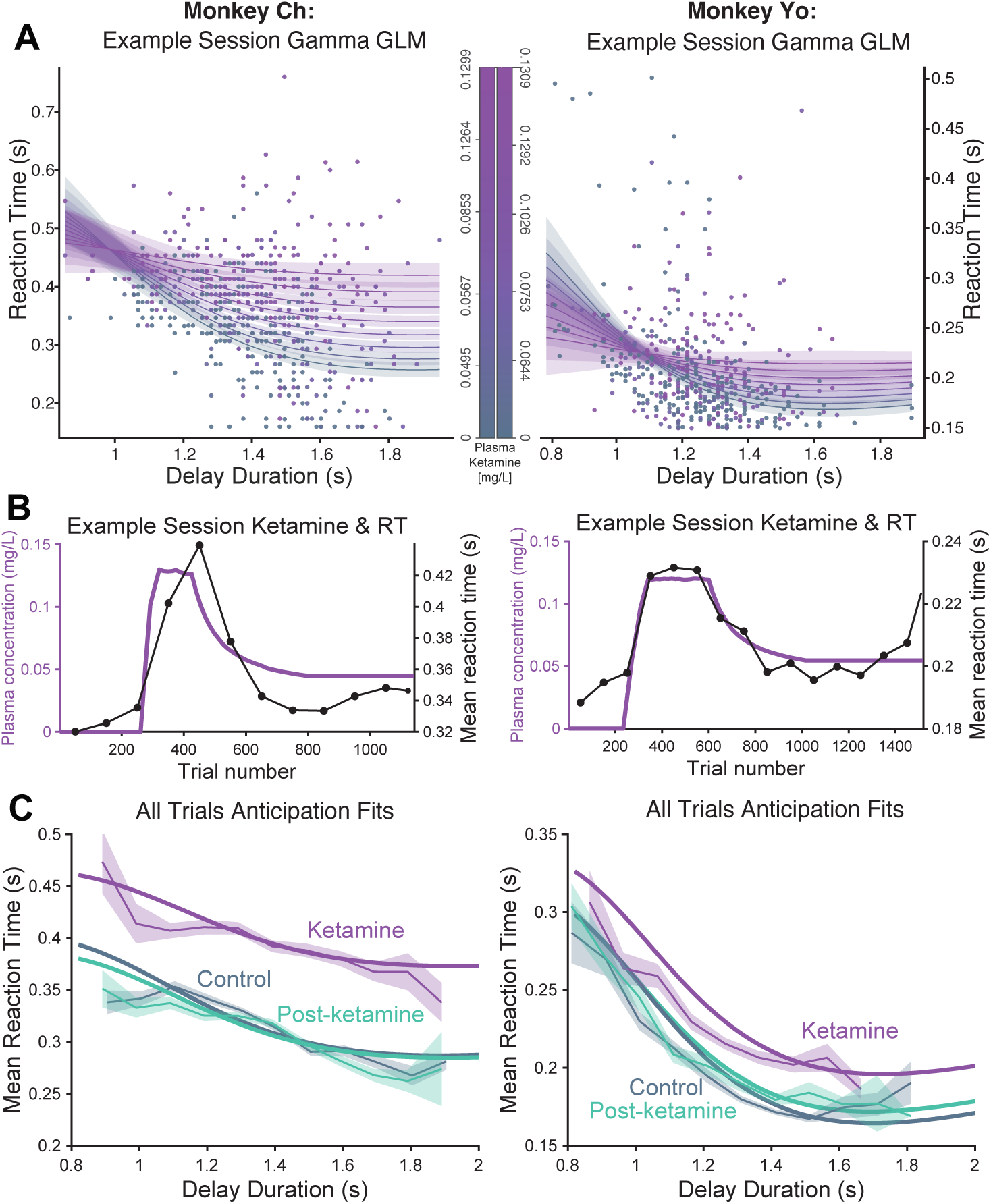
Anticipation across ketamine concentrations. **(A)** Gamma GLM fits to RTs (lines) overlaid on individual trial RTs (dots) for one example session in each monkey. The fits are shown across the full range of plasma ketamine concentrations (colorbar), with each dot colored by the ketamine concentration on that trial. **(B)** Mean RT across 100-trial bins (black) along with estimated plasma concentration from the pharmacokinetic model of ketamine for an example session (Equation 2). **(C)** Mean RT (±SE) as a function of delay duration across all trials from each monkey, under control (gray), steady-state ketamine (purple), and post-ketamine (green) conditions. Solid curves are fits of the anticipation models in Figure 1B.

**Figure S3:**
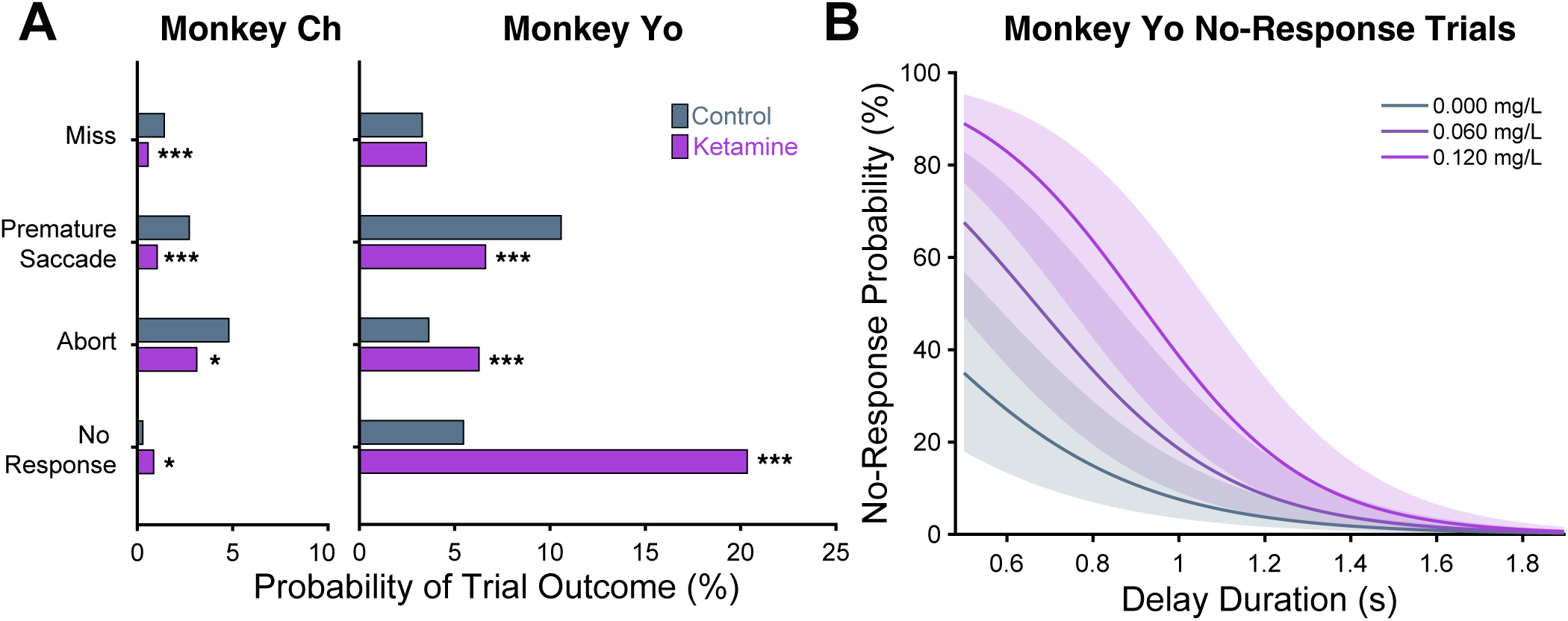
Change in trial outcomes under ketamine. **(A)** Probability of each type of unrewarded trial (see Methods) under control and ketamine conditions. Significance of control versus ketamine mixed model effects in each category is indicated with ∗ : *p* < 0.05, *∗ ∗ ∗* : *p* < 0.0001. **(B)** The probability of no-response trials in Monkey Yo as a function of delay, for control condition and two ketamine concentrations (Equation 8). No-responses were more likely at shorter delays (*β*_2_ = *−*3.74, *p* < 10^−48^), and this bias toward shorter delays grew with ketamine concentration (*β*_3_ = *−*11.3, *p* = 0.007).

**Figure S4:**
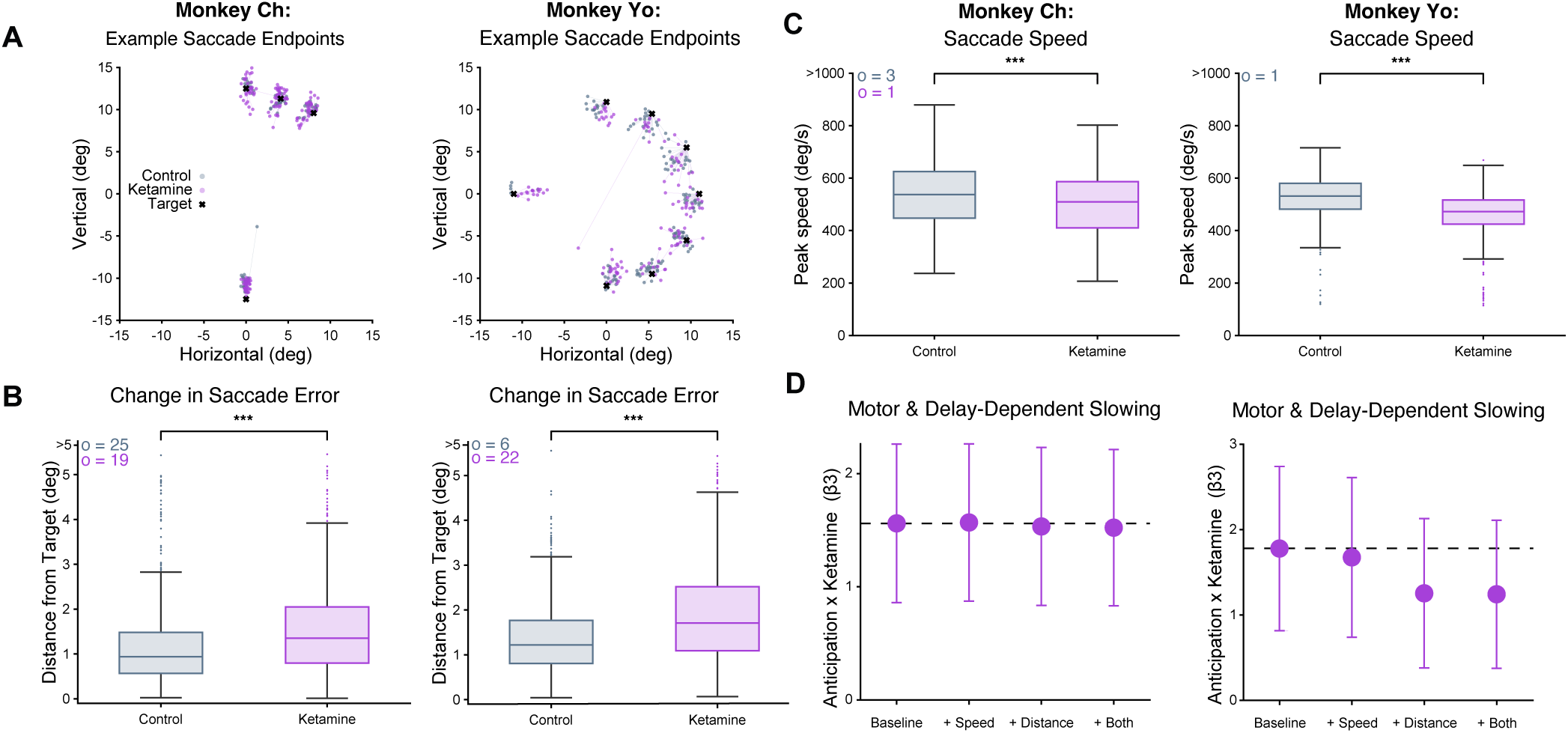
Saccade velocity and accuracy change under ketamine. **(A)** Saccade endpoints for correct and missed trials across one example session for Monkey Ch and Monkey Yo. Pre-ketamine control trials are indicated by dots in grey and steady-state ketamine trials are in purple. Target positions are indicated by black Xs, with lines connecting eye positions to the intended target. **(B)** Change of distance from target to endpoint eye-position under ketamine for each monkey, including both correct and missed trials (: *p* < 0.001, see Methods). Boxplots indicate median distance and the first and third quartiles, with dots indicating outliers. Counts of outliers (o) *>* 5 deg are indicated in the upper left. **(C)** Boxplots of saccade velocity under control and ketamine conditions for each monkey, with the same conventions as in *B*. **(D)** Delay-dependent effect of ketamine (*β*_3_) for the gamma GLM with and without speed and endpoint error predictors (Equation 10), with 95% confidence intervals. Baseline estimate is indicated by the dashed line. *β*_3_ remained significant in all models for both monkeys.

**Figure S5:**
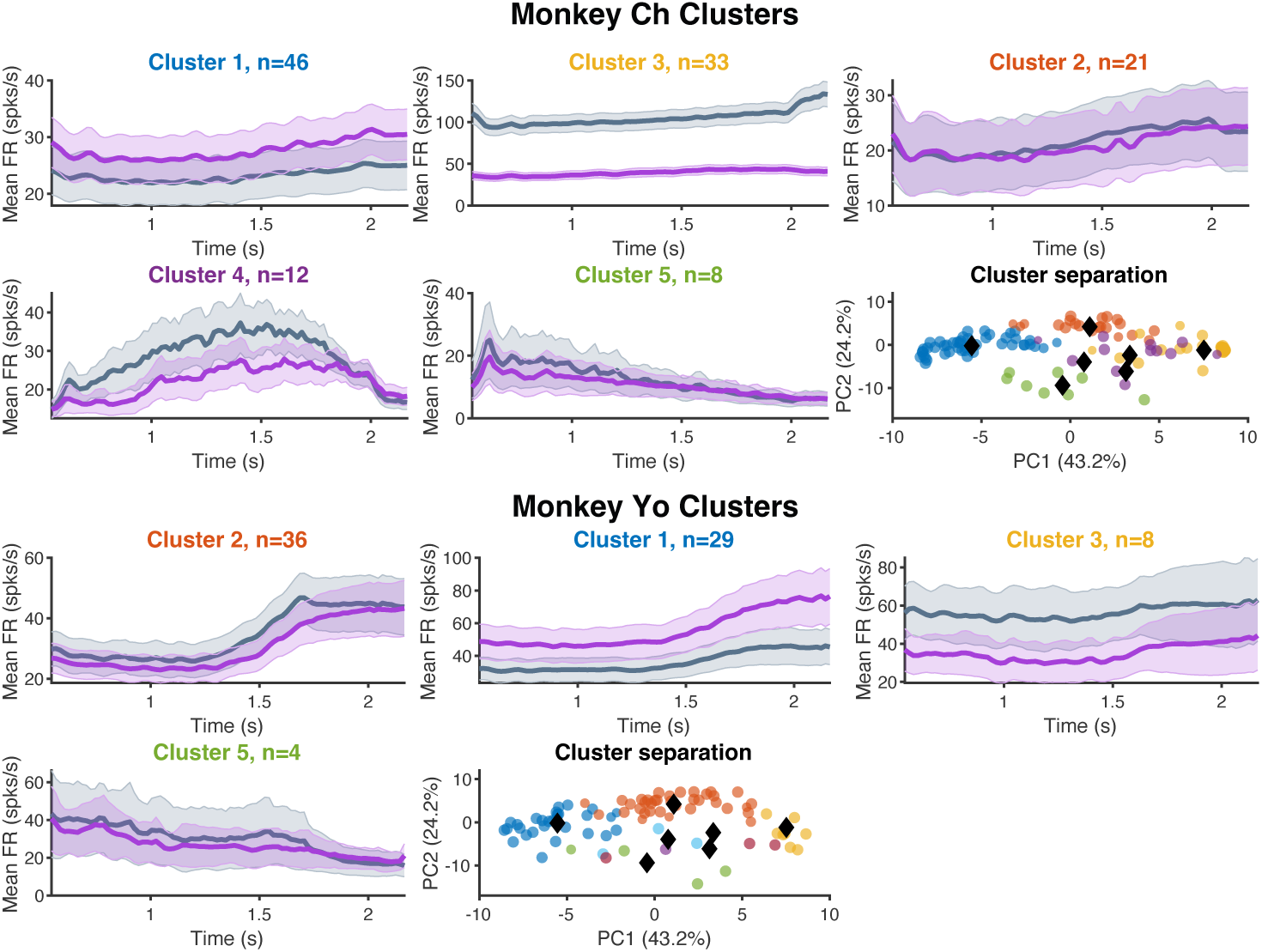
GMM clusters separated by monkey. Mean delay kernel activity (±SE) for clusters identified through GMM on the combined population (Figure 3), shown separately for each monkey. For clarity, only clusters with more than 3 neurons are shown, ordered by descending number of neurons within each monkey. Activity is shown under control (gray) and steady-state ketamine (purple) conditions. Cluster positions in PCA space are shown at bottom right for each monkey.

**Figure S6:**
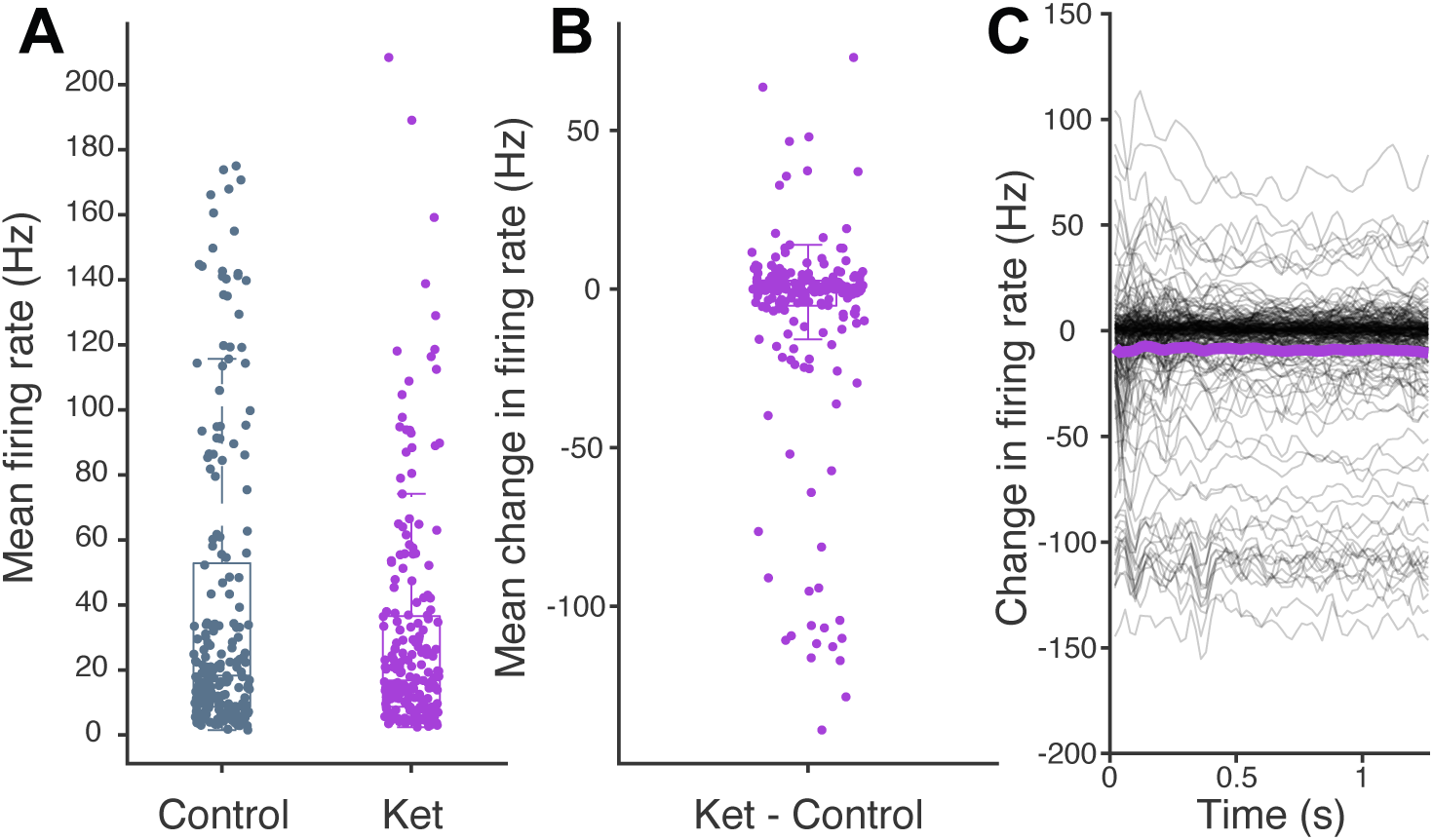
Firing rate changes under Ketamine. Effect of ketamine on delay-period firing rate is shown across the recorded population, pooled from the two monkeys. **(A)** Mean delay-period firing rate for each neuron under pre-ketamine control and steady-state ketamine. **(B)** Distribution of per-neuron changes in mean firing rate (ketamine–control) is centered near zero, with neurons showing both increases and decreases under ketamine. **(C)** Change in firing rate across the delay period for each neuron.

**Figure S7:**
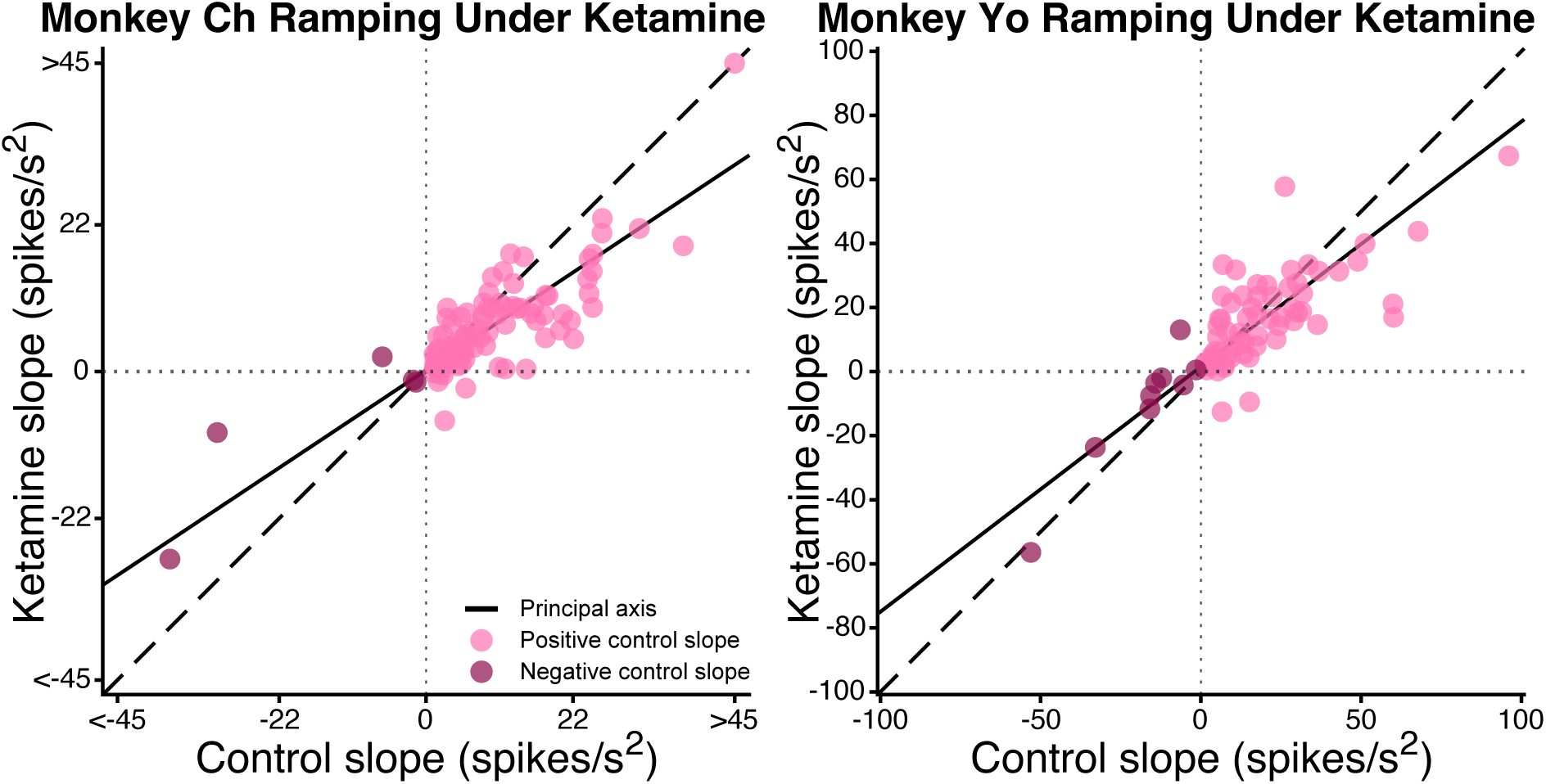
Effect of ketamine on slope of delay-period firing rates. Slope of the delay-period firing rate under steady-state ketamine versus control is plotted separately for each monkey. Points represent individual neurons and are colored by the sign of the pre-ketamine control slope (upward or downward ramping). The principal axis is shown as a solid line and the unity line is dashed. This is the raw firing rate counterpart to slopes computed from delay kernel activity shown in Figure 4E-F.

**Figure S8:**
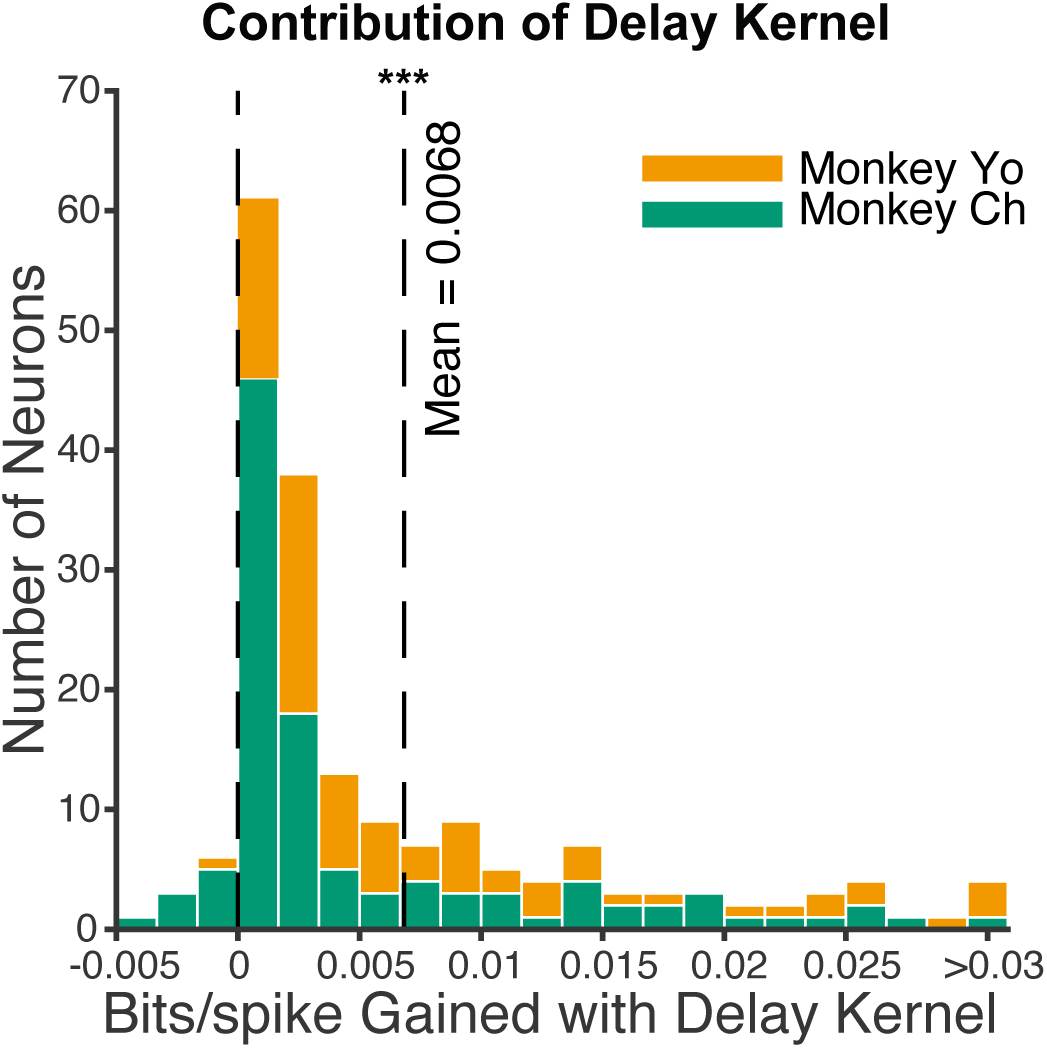
Delay kernel improves spike prediction in ketamine model. Cross-validated model performance (bits/spike) for the ketamine-parameterized model is compared between the full model (with delay kernel) and a reduced model (without delay kernel) for each neuron, in the two monkeys. Similar to Figure 3B, which showed the same comparison for control data, values greater than zero indicate neurons for which inclusion of a delay-period kernel improved spike prediction in the ketamine model.

**Figure S9:**
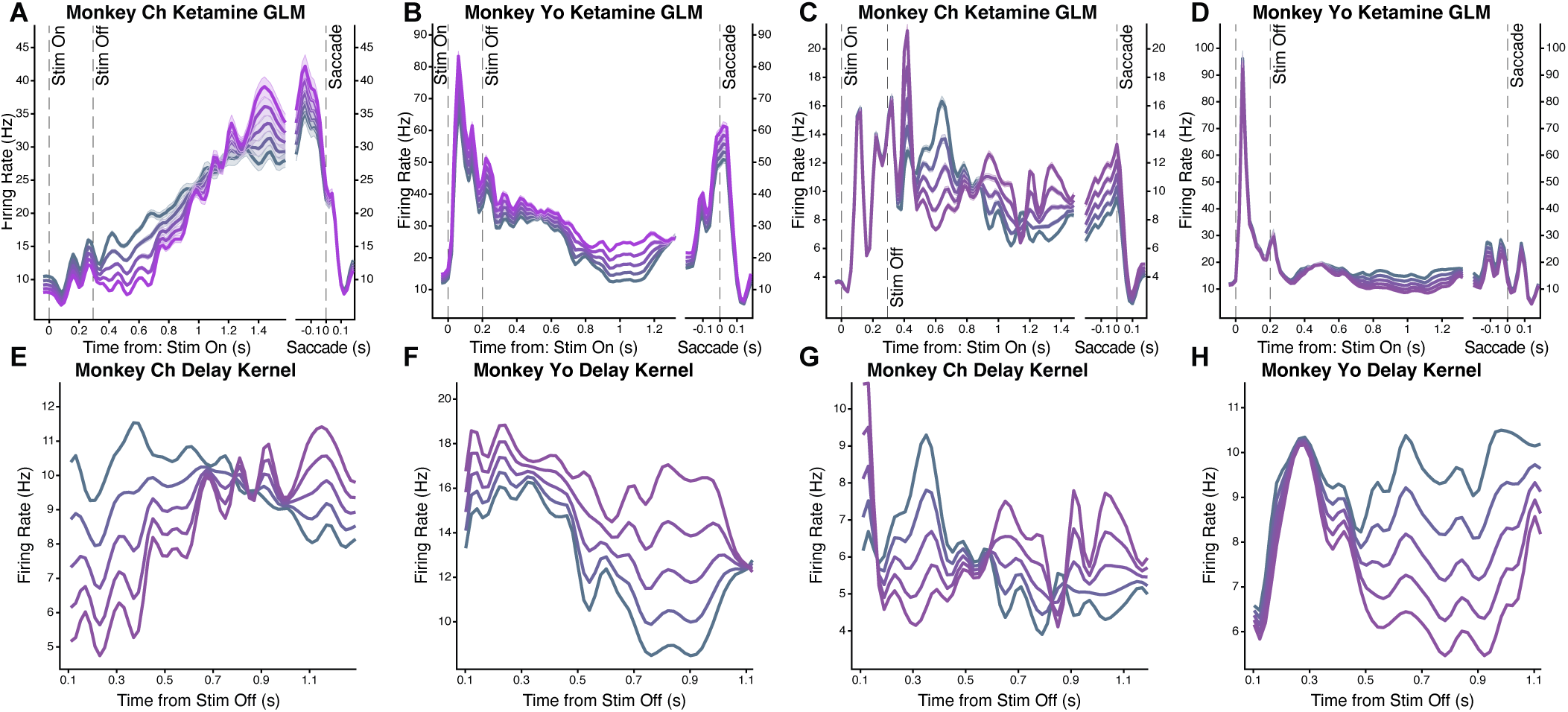
Predictions from ketamine-parameterized model. **(A-D)** Predictions for example neurons from the ketamine-parameterized model, two from each monkey. Stimulus onset and saccade-aligned epochs are shown. **(E-H)** Isolated delay kernel activity for the same neurons (matched by column) by time from stimulus offset. Delay kernels began 50 ms after stimulus offset and overlapped with the sensory kernel for an additional 50 ms.

**Figure S10:**
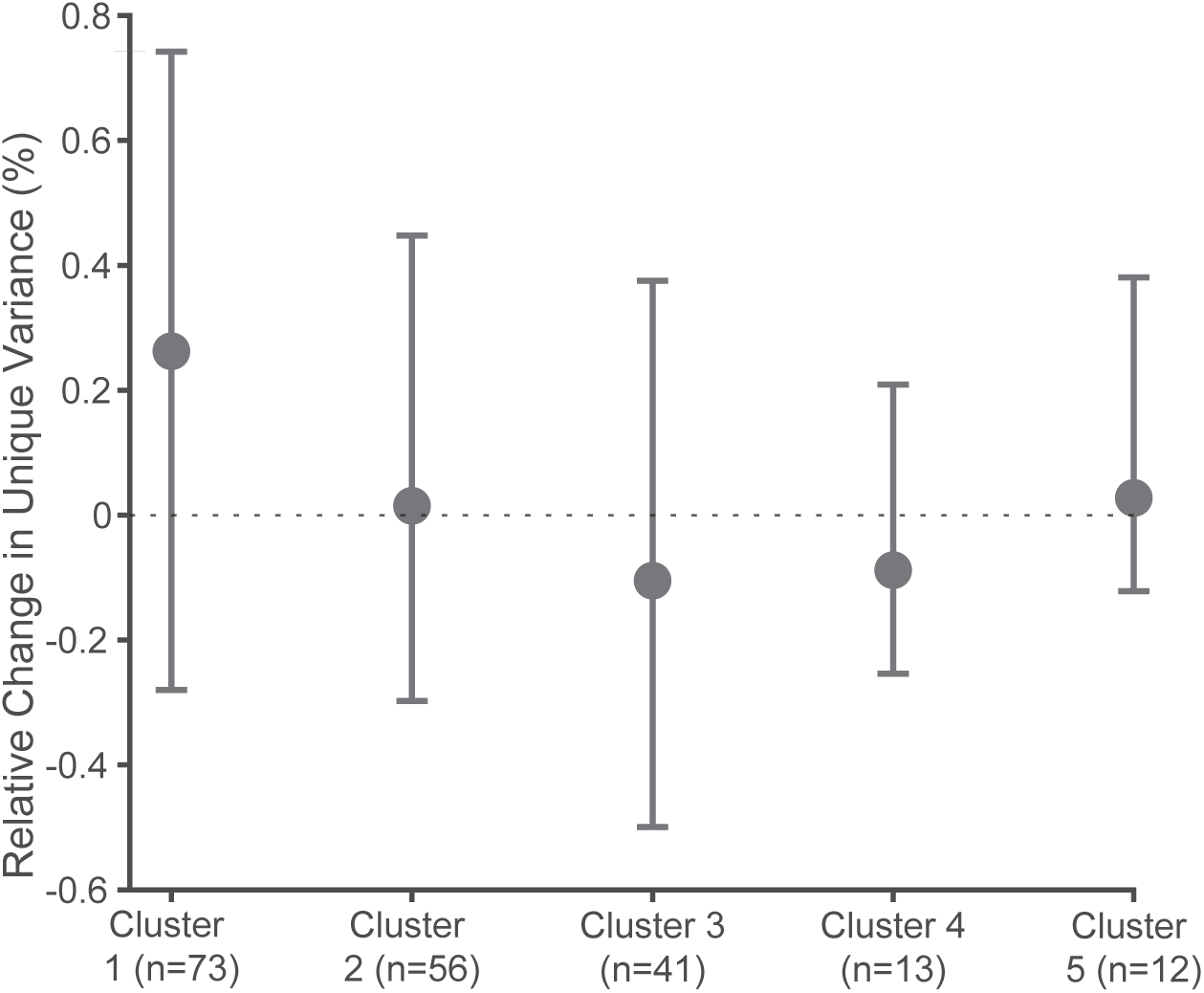
Behavioral Time Axis Variance With Cluster Removal. Change in the unique variance explained by the behavioral time axis when each cluster is removed prior to dPCA fitting, relative to removal of a size-matched random set of neurons. Removing any single cluster, including the upward ramping clusters (#1 and #2) did not significantly reduce the axis variance, indicating that no single cluster is necessary for the axis.

**Figure S11:**
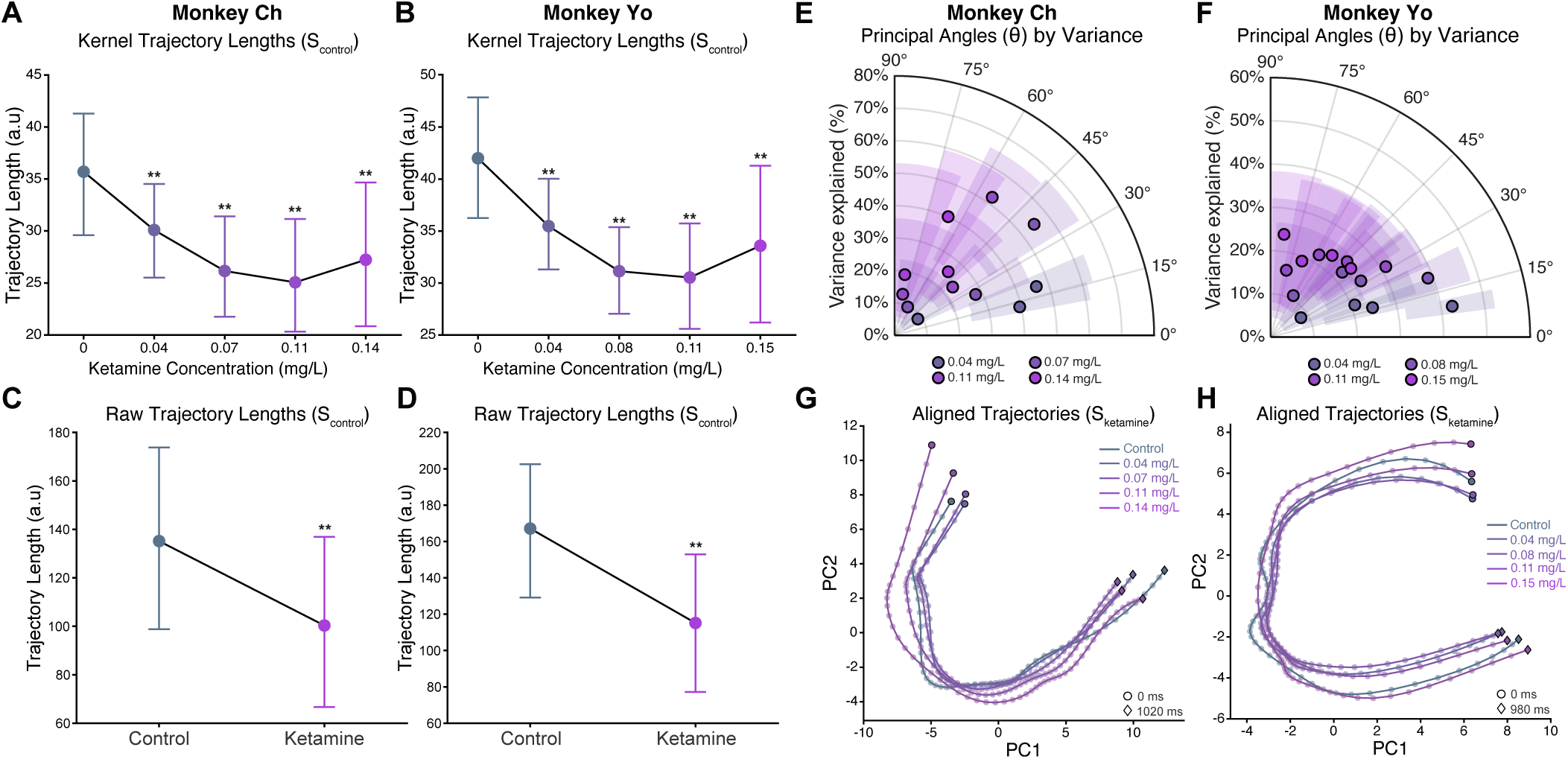
Trajectory lengths under ketamine. **(A-B)** Population trajectory length at a range of ketamine plasma concentrations predicted from GLM delay kernels, projected into *S_control_*. Projected trajectories decreased in length under ketamine in both monkeys, and bootstrap 95% CIs are indicated by bars along with markers of significance (: *p* < 0.01). **(C-D)** Population trajectory length at baseline versus steady-state ketamine, computed from raw delay-period firing rates and projected into the control subspace (*S_control_*). **(E-F)** Principal angles between *S_control_* and *S_ketamine_* for a range of ketamine concentrations, with 0° being alignment of the associated principal vectors. Radial position indicates the percent of each ketamine condition’s total variance captured by the associated principal vector. Colors indicate ketamine concentration. Points show bootstrap means and shaded areas indicate 95% CIs for angle and variance explained. **(G-H)** Trajectories derived from delay kernels for a range of ketamine concentrations, calculated separately for each concentration (*S_ketamine_*) and aligned using Procrustes analysis. In contrast to projections onto *S_control_* which are distorted by subspace rotation, these trajectories demonstrate conservation of shape across concentrations.

